# Independent adoptions of a set of proteins found in the matrix of the mineralized shell-like eggcase of Argonaut octopuses

**DOI:** 10.1101/2021.07.10.451900

**Authors:** Davin H. E. Setiamarga, Kazuki Hirota, Risa Ikai, Seiji Imoto, Noriyoshi Sato, Hiroki Ono, Yukinobu Isowa, Hiroshi Yonemitsu, Takenori Sasaki, Masa-aki Yoshida

## Abstract

The Argonaut octopus, commonly called the paper nautilus, has a spiral-coiled shell-like eggcase. As the main characteristics, the eggcase has no internal septum, is composed entirely of calcite with chitosan being the main polycarbonate and is reportedly formed by organic materials secreted from the membranes of the arms. Meanwhile, the biomineralized external “true” shells of the Mollusks, which includes the Cephalopods, are secreted from the mantle tissue. Therefore, the histological origin of the two shells is completely different. The question of how the Argonauts, which phylogenetically diverged from the completely shell-less octopuses, could form a converging shell-like external structure has thus intrigued biologists for a long time. To answer this question, we performed a multi-omics analysis of the transcriptome and proteome of the two congeneric Argonaut species, *Argonauta argo* and *A. hians*. Our result indicates that the shell-like eggcase is not a homolog of the shell, even at the protein level, because the Argonauts apparently recruited a different set of protein repertoires to as eggcase matrix proteins (EcMPs). However, we also found the homologs of three shell matrix proteins (SMPs) of the Conchiferan Mollusks, Pif-like, SOD, and TRX, in the eggcase matrix. The proteins were also found in the only surviving shelled Cephalopods, the nautiloid *Nautilus pompilius*. Phylogenetic analysis revealed that homologous genes of the Conchiferan SMPs and EcMPs were found in the draft genome of shell-less octopuses. Our result reported here thus suggests that the SMP-coding genes are conserved in both shelled and shell-less Cephalopods. Meanwhile, the Argonauts adopted some of the SMP-coding genes and other non-SMP-coding genes, to form a convergent, non-homologous biomineralized external structure, the eggcase, which is autapomorphic to the group.

## Introduction

The presence of a biomineralized external shell is one of the defining characters of Conchifera (Cowen 2009), a subphylum of Mollusca composed of five shelled molluscan orders (Monoplacophorans, Cephalopods, Scaphopods, Gastropods, and Bivalves) (Kocot et al. 2010; Kocot et al. 2020; Smith et al. 2011). Calcium carbonate-based biominerals (e.g., aragonite, calcite) are known to be the main component of Molluscan shells (Spann et al. 2010; Frenzel and Harper 2011). Interestingly, however, while the Cephalopods includes famous extinct members with univalve shells (e.g., ammonites, ortocherids, and belemnites), almost all of the ca. 800 species of extant cephalopods had evolutionarily internalized, reduced, or completely lost their shells (Kröger et al. 2011). Only the single surviving family of the Nautiloidea, the Nautiliids, still retain their true calcified external shell (e.g., Setiamarga et al. 2020). 300 of the extant cephalopod species are members of the order Octopodiformes, which include octopuses and vampire squids (Sanchez et al. 2016). Because of the lack of stable morphological characters, the systematics of this group has been deemed difficult (Strugnell et al. 2011). However, various recent molecular phylogenetics studies have consistently shown that members of this group can be classified into two suborders, Cirrata (cirrate octopuses) and Incirrata, and that the incirrate octopuses can be further grouped into six families: Amphitretidae, Alloposidae, Argonautidae, Ocythoidae, Tremoctopodidae, and Octopodidae, with the first five families, pelagic (Strugnell et al. 2006; Strugnell et al. 2011; Sanchez et al. 2016; Chiu et al. 2018). Despite being a member of the Conchiferans, all extant members of Octopodiformes are known to have lost their calcified external shell (Nishiguchi and Royal 2008; Kröger et al. 2011; Ponder et al. 2019).

Within the five families of the incirrate octopuses, only the argonauts (Argonautidae) exceptionally have a calcified external shell-like structure (an “eggcase”) which resembles the coiled shell of the Nautilids and Ammonoids (or, like a hollyhock leaf when folded in the center), although it lacks true Conchiferan shell microstructures and very brittle (Naef, 1923; Wolfe et al. 2012) (Fig. 1AB). The shell-like structure, though, is only seen in females, is thought to play a role in protecting the eggs laid inside, as well as taking in air for buoyancy (Finn and Norman 2010). In the past, morphological studies have established Argonautidae as its own monophyletic group composed of four described extant species (*Argonauta argo*, *A*. *hians*, *A. cornuta*, and *A*. *boettgeri*), based on the presence of this shell-like eggcase. However, recent molecular reassessments have confidently grouped Argonautidae together with the shelless blanket octopuses Tremoctopodidae (Strugnell et al. 2006; Hirota et al. 2021). The phylogeny thus seems to support the suggestion that the shell (eggcase) of the argonauts is not a true shell and thus not a homolog of the calcified shell of Conchiferan Mollusks. The eggcase is thus most likely acquired secondarily from a shell-less Octopodiform ancestor (Wolfe et al. 2012) they shared with the tremoctopodids, and thus have evolved as an evolutionary innovation (a synapomorphy) of Argonautidae (Naef, 1923).

It is very curious how the argonauts reacquired a biomineralized structure independently, from a putative ancestral octopod which has lost its external calcified tissues. In general, Molluscan shell is basically formed through the additive growth of biomineralized tissues through continuous secretion from the mantle tissue under a genetic control (Carter 1990). The genetic control regulates the production of the organic part of the shell matrix, causing the minerals to have distinct layouts called the microstructure, in a process called the “organic matrix-mediated mineralization” (Lowenstam 1981; Weiner et al. 1983; Takeuchi et al. 2008; Marin et al. 2012). Scales (2015) mentioned that in the 19th century, Villeprez observed that the first arms were involved in the shells repairing/formation in *Argonauta argo*. Thus ontogenetically, the true shells of Conchiferans (including those of shelled Cephalopods) differ from the eggcase of the Argonauts, but only because the putative organic compound secreting tissue is located in the first arms, rather than the mantle tissue. Besides that, at the first glance, the eggcase of the Argonauts appears morphologically similar to the biomineralized external shell of the Conchiferans, but they are microstructurally different (Oudot et al. 2020b). The eggcase of the Argonauts contains only calcite crystals while the outer shell of molluscs contains mainly aragonite and calcite (Revelle and Fairbridge 1957; Mitchell et al. 1994; Nixon and Young 2003; Saul and Stadum 2005). The microstructure of the eggcase, where the core is sandwiched between two fiber layers, is also unique to the Argonauts (Oudot et al. 2020b). However, it is without doubt that the Argonauts also adopt the “organic matrix-mediated mineralization” to form its shell, albeit using a different secretive tissue. This curiosity becomes further unclear, because the molecular evolution of the genetic components involved in the eggcase formation, were not yet properly studied. This is probably because fresh Argonauts samples needed for molecular studies are difficult to obtain, and living individuals are difficult to keep in aquaria.

Recent development in genome, transcriptome, and proteome sequencings, however, has allowed for large-scale molecular dissection (e.g., Zhang et al. 2012; Mann et al. 2012; Marie et al. 2012; Miyamoto et al. 2013; Arivalagan et al. 2017; Zhao et al. 2018; Setiamarga et al. 2020) and identification (e.g., Miyamoto et al. 1996; Sudo et al. 1997; Tsukamoto et al. 2004) of major proteins in the shell of the Conchiferans (e.g., Miyamoto et al. 1996; Sudo et al. 1997; Tsukamoto et al. 2004). Many of these proteins are present in trace amounts inside the shell and are therefore referred to as shell matrix proteins (SMPs), which are also a part of the organic component of the shell. Despite their low abundance, the SMPs play an essential role in shell formation and structural maintenance, including calcium carbonate nucleation, crystal growth, and selection of calcium carbonate polymorphs (Addadi et al. 2006; Marin et al. 2008). Interestingly, many of these proteins mostly contain well-characterized domains (repetitive low-complexity domains (RLCDs), peroxidases, carbonic anhydrases (CAs), chitinases, acidic calcium-binding proteins, and protease inhibitors) and thus are homologs of some non-shell matrix known proteins (Marie et al. 2012). Other SMPs include acidic proteins probably involved in CaCO3 crystallization, such as Chitin-binding proteins and ECM-related proteins (Pif, BMSP, EGF-ZP, SPARC) (Dyachuk 2018, Feng et al. 2017). The finding of highly conserved domains across the Conchiferanss indicates that SMPs have existed as ancestral genes for a long time (Kocot et al. 2016; Setiamarga et al. 2020). However, molecular evolution studies have also shown that they might have been co-opted independently as SMPs in different molluscan lineages. For example, independent recruitments of CAs were thought to happen in *C. gigas* and *P. fucata* after their divergence (Zhao et al. 2020). Besides that, SMPs classified as RLCDs have undergone a rapid parallel evolution in the Mollusks (McDougall et al. 2013; Jackson et al. 2010). Meanwhile, Glycine-rich RLCDs, shematrin, and KRMP gene families found in the pearl oysters have undergone multiple duplications, extensive sequence divergence, and shuffling of various motifs (Takeuchi et al. 2016; Kenny et al. 2020; Zhang et al. 2012). The availability of these bulk data of SMPs and the molecular evolution of some of the SMP-coding genes has allowed researchers to build a core set of Conchiferan SMPs, which will be useful for comparative analyses. Recently, we conducted a study on the basal Cephalopods, the “living fossil” *Nautilus pompilius*, which still retains its biomineralized external shell and found a set of core Conchiferan SMPs and their protein domains in the *Nautilus* shell (Setiamarga et al. 2020). The results of these studies could then be used to elucidate the molecular evolution behind the phenotypic convergence of a morphological structure, that is, the reacquisition of a shell-like structure in the Argonauts.

At the genetic level, it was unclear whether shell loss in Octopodiformes involved gene losses or not, while the reacquisition of a calcified eggcase in the Argonauts was a result of reacquisitions of genes with similar functions (gene reactivations), or novel recruitments. In general, loss-of-function in genes would cause the genes to become pseudogenes, and eventually, be removed from the genome. However, if genes related to shell formation were completely pruned out, we then wonder how the ability to form shells could evolve from an octopus without any functioning biomineralization genetic tool. The first hypothesis is that the shell-forming gene was lost in the ancestors of the octopus, and other genes took over the role of forming the outer shell in the argonauts to replace the lost gene (novel recruitments). In this case, the SMPs detected in the Argonauts are expected to be very different from the standard gene set of the Conchiferans (Setiamarga et al. 2020). On the other hand, there is a known case in which degeneration of the shell results in small changes at the genetic level. The sea hare (*Aplysia californica*) has a weakly calcified inner shell that has completely lost its original function of protecting soft tissues. However, a subset of the protein repertoire is conserved in the sea hare shell, including Tyrosinases, Kunitz Protease Inhibitors, and RLCD-containing Proteins, which exhibit characteristics similar to those of a fully functional outer shell (Marin et al. 2018). This thus leads to the second hypothesis, where the shell-forming genes are somehow conserved in the octopus (probably because they are multifunctional), but have lost their specific functions in shell formation. In this case, although we still cannot say that the eggcase of the Argonauts is homologous to the shell of the Conchiferans, it might show a deep homology between the shell-like eggcase of the Argonauts and the actual shell of the Conchiferans (Shubin et al. 2009; Hall 2012; McCune and Schimenti 2012; Tschopp and Tabin 2017).

In order to assess these hypotheses, we performed a multi-omics analysis of the transcriptome and proteome in two species of Argonauts, *Argonauta argo* and *A. hians*, and conducted bioinformatics comparative analyses among the Argonauts and some Conchiferans for which the SMP data is already available, including those of *Octopus bimaculatus*. The result of our analyses, which in general can be summarized as (1) no Conchiferan homologous SMP-coding gene was present in the eggcase matrix of the Argonauts, (2) Conchiferan SMP homologs (or homologous domain) were found in the genome of the shell-less octopus, *Octopus bimaculatus*, and (3) the proteins found in the eggcase matrix of the Argonauts are homologous with non-shell matrix-forming genes in other animals, can thus be said to support the second hypothesis.

## Materials and Methods

### Sample collection

*Argonauta argo*, *A*. *hians*, and *A*. *boettgeri* are known to be cosmopolitan and distributed in tropical and subtropical waters worldwide. However, *A. argo* and *A*. *hians* drift in warm currents to the Sea of Japan to undergo a mortality migration, dying under low temperatures in winter. Shimane Prefecture in Japan is located on the Sea of Japan along the path of the warm current (called the Tsushima current), and it is possible to obtain living or fresh specimens of *A. argo* and *A. hians* by fixed nets from June to August (Sakurai and Kono 2010, Fig. 1C). Thus, this has allowed us to obtain enough specimens to conduct our study.

**Fig. 1.**
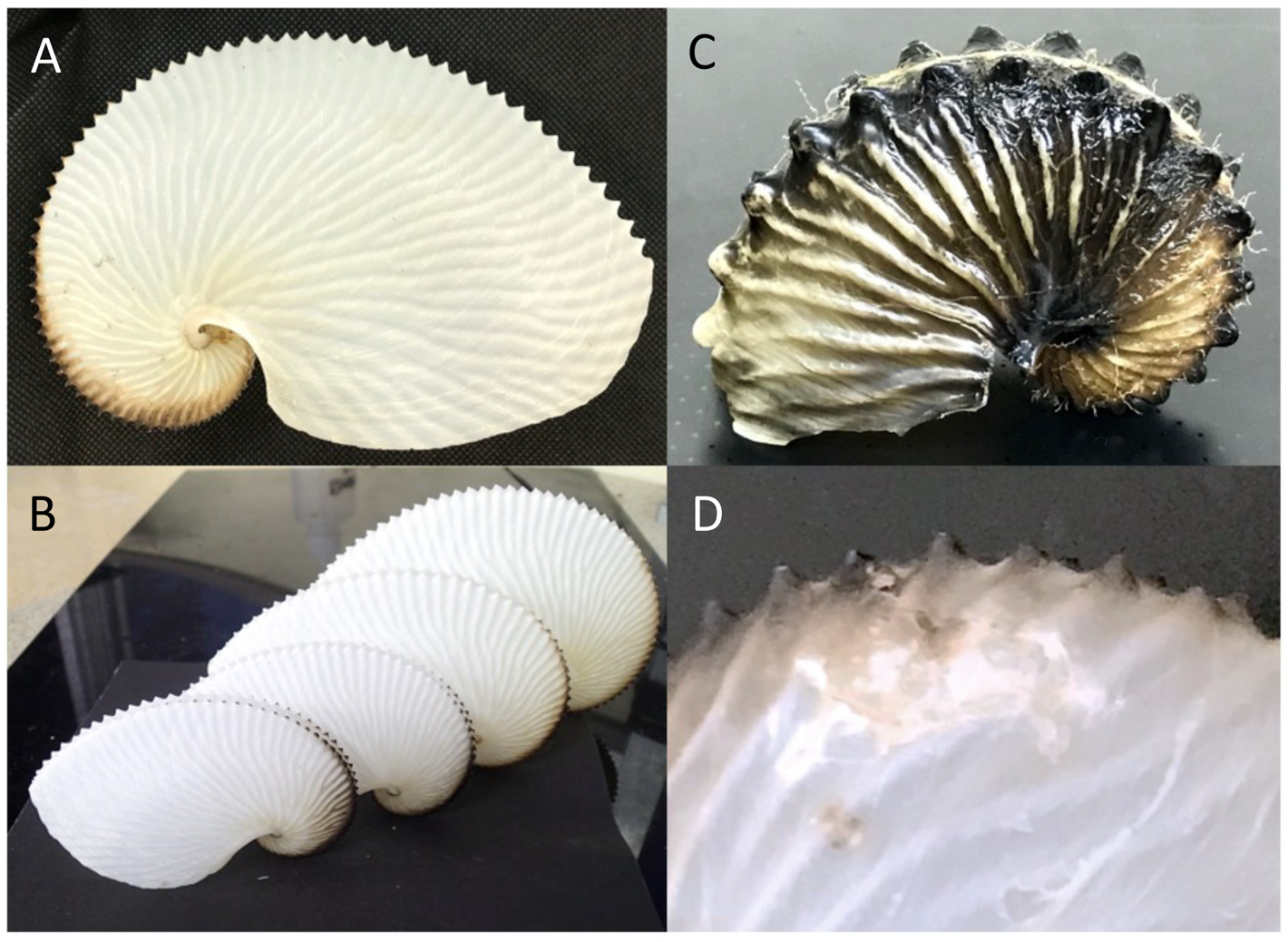
The eggcases (“shells”) of Argonaut octopuses. A. The shell-like eggcase of *Argonauta argo*. B. The shell-like eggcase of *Argonauta argo* from several individuals of different sizes, most likely because of individual growth in body size. C. The shell-like eggcase of *Argonauta hians*. D. The shell-like eggcase of *Argonauta argo* showing a trace of a possible shell repairment.

Fresh specimens of *A. argo* and *A. hians* were provided by the fishermen from the bycatch caught by the fixed nets set along the coast in Oki Island Town, Shimane Prefecture, Japan (36°17’20.6"N 133°12’46.4"E). Pieces of the mantle, membrane of the first arm, and tips of the second arm were obtained from a single individual. For *A. argo*, the eyes, hearts, and gill hearts were also sampled from different individuals. Total RNA was extracted from the tissue samples using Trizol followed by the RNeasy mini kit (Qiagen) with an on-column DNaseI treatment. RNA aliquots were stored in –80°C until further transcriptome analyses.

### Transcriptome analyses

Transcriptome analysis of the arms and mantle was outsourced to Macrogen Japan, Inc, using Illumina HiSeq 2000, under the 2 × 100 bp paired-end platform (PE100). Total RNA from of the eye, heart, and gill heart were sequenced on an Illumina HiSeq 2000 and 2500 at the National Institute of Genetics, Japan (NIG) under the support of the Japanese Government through the Platform for Advanced Genome Science (PAGS) project. Raw read sequence data will be available in the DNA Data Bank of Japan (DDBJ). We are willing to share our raw data before the publication of the original paper on the assumption that it will be done as a collaborative research.

Obtained raw reads from the six libraries were combined for *A*. *argo*, and then assembled using the Trinity assembly v.2.8.5 on the supercomputing system of the NIG. Similarly, the three libraries were assembled together for *A. hians*. Six frame translations of the Trinity assembly contigs were obtained by using a custom perl script and used for the proteome database. For the expression profiling, the FASTQ reads are aligned to the Trinity contigs using kallisto (version 0.44.0) and salmon (version 0.14.1) (Bray et al. 2016; Patro et al. 2017). The gene expression matrix of transcripts per kilobase million (TPM) was summerised with R/tximport (Soneson et al. 2015). Probable GPI anchor site was predicted using PredGPI (Pierleoni et al. 2008).

### Proteome sequencing of eggcase matrix proteins of *A*. *argo* and *A. hians*

Several eggcasses obtained from multiple individuals of the two argonaut species were used in proteome sequencings. Several shells of one species were first shattered into pieces using a hammer, and then cleaned from any possible external organic contaminants through incubation in a 2M NaOH overnight. Cleaned shell pieces were then ground into powder, and then slowly decalcified using 0.5 M EDTA as the chelating agent, at 4°C for 3 days. Total hydrophilic proteins of the shell were extracted and purified using the 3 kDa Amicon Ultra Centrifugal Filter Unit, with the hydrophilic fraction obtained as eluate in MilliQ water, which then was stored at −80°C until further analyses. Hydrophobic proteins were obtained as pellets. After a thorough cleansing using MilliQ water, the pellet was eluted in a 9M urea solution, and then further cleansed several times using a standard acetone precipitation method. Collected final precipitate was also stored at −80°C until further analyses. The presence of proteins in both fractions was confirmed using standard SDS-PAGE and silver staining methods.

Protein sequencing was outsourced to Medical ProteoScope Co., Ltd. (Yokohama, Japan), under the following protocols: After a short deployment of SDS-PAGE, samples were extracted from the gel and subsequently were partially digested into short peptides with trypsin, following Shevchenko et al. (1996). Peptide samples were sequenced using a DiNa nanoLC Liquid system (KYA Technologies, Tokyo, Japan) and a LTQ Orbitrap mass spectrometer (Thermo Fisher Scientific). The resulting MS/MS spectra data was searched against the protein sequence database using a software search engine of Mascot (ver. 2.6.2) (Matrix Science), with the index of False Discovery Rate (FDR) set to 1%. A custom protein sequence database containing the six-frame translated transcriptome sequences and 116 sequences of possible contaminants collected from The Global Proteome Machine Organization (http://www.thegpm.org/crap/index.html) was assembled and used for the searches.

### Annotations of the eggcase matrix protein sequences of *A*. *argo* and *A. hians*

Sequence annotation of identified transcripts was performed by conducting reciprocal local BLASTp and BLASTx searches on three databases: (1) the nr database of Genbank, and (2) a database of Conchiferan shell matrix proteins we assembled from data published until March, 2020 (Setiamarga et al, 2020), and (3) *Octopus bimaculatus* genome data. Protein domains were predicted using the SMART domain search online tool (http://smart.embl-heidelberg.de/; Letunic et al. 2021) under the “Genomic” mode. The result was confirmed using the online versions of InterProScan (https://www.ebi.ac.uk/interpro/search/sequence/; Jones et al. 2014) under the default settings on the browser, and Prosite (https://prosite.expasy.org/; De Castro et al. 2006) under the Quick Scan mode. Signal peptides were predicted using the online tool SignalP (http://www.cbs.dtu.dk/services/SignalP/; Petersen et al. 2011) under the default settings on the browser. Predicted domains were then visualized using an R script. Only the result as indicated by SMART and SignalP was shown in Figure 2A, 2B.

**Fig. 2.**
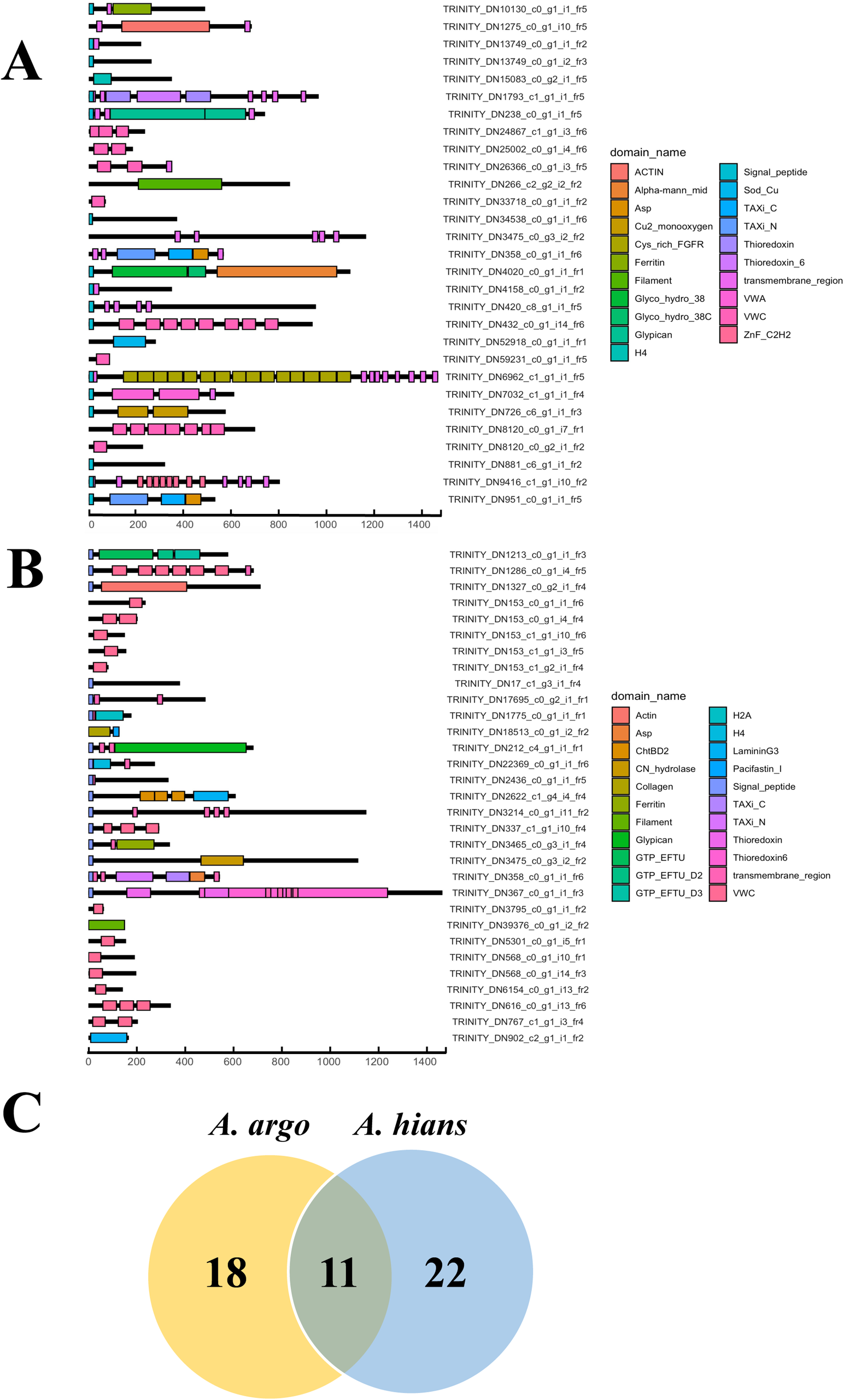
Eggcase matrix proteins of the Argonauts, identified through the multi-omics approach in this study. A. The domain structures of eggcase matrix proteins (EcMPs) of *Argonauta argo* B. The domain structures of the eggcase matrix proteins (EcMPs) of *Argonauta hians* C. A Venn diagram showing the number of shared EcMPs between the two congeneric species of the Argonauts

### Comparative analysis of various Conchiferan Shell Matrix Proteins

We first conducted reciprocal local BLASTp searches using two search settings (Search Settings 1 = e-value <1e-50 and threshold ≥40%; Search Settings 2 = e-value <1e-5), of the annotated 29 *A. argo* and 33 *A. hians* eggcase matrix protein sequences (EcMPs), in order to see which proteins are shared between the two species. The result was plotted as a Venn diagram in Fig. 2C. Meanwhile, a Circos chart showing the shared proteins, with the sequences arranged based on their emPAI data and sequence lengths, is presented in Supplementary Figure 1. Circos charts in this study were drawn using the software Circos-0.69-9 (http://circos.ca/).

Next, we also conducted local BLASTp searches of the annotated EcMPs of *A. argo* and *A. hians*, against the shell matrix protein data of *Nautilus pompilius* (Setiamarga et al. 2020), and the genome data of *Octopus bimaculatus* (Albertin et al. 2015). From these searches, we expected to obtain information about possible conservation of the SMPs and EcMPs among shelled and shell-less cephalopods. However, because the divergence of all Cephalopods happened very ancient (Mid-Silurian, ca. 432 MYA; Sanchez et al, 2016; Tanner et al. 2017), we only showed and further discussed the result of Search Setting 2 in the main manuscript (Fig. 3).

**Fig. 3.**
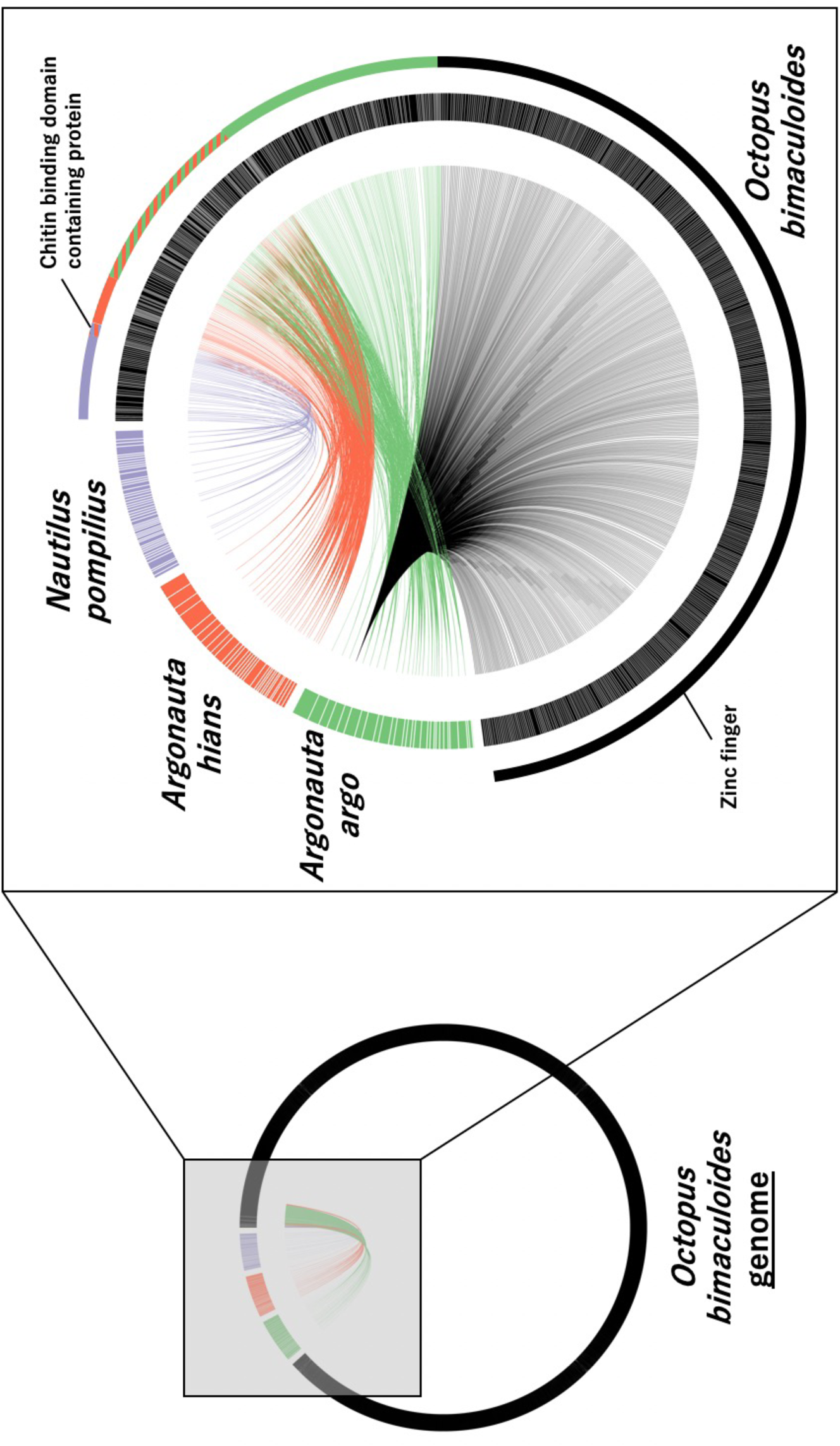
Comparison of Cephalopod Shell Matrix Proteins (SMPs). Reciprocal BLASTx searches among the different species of Cephalopods, in order to see if the Argonauts use the same SMPs as the shelled basal Cephalopods, *Nautilus pompilius*, which were shown to retain many of the Conchiferan SMPs in its shell (Setiamarga et al., 2020). The result reciprocal BLASTx and genomic searches were also conducted among the genome data of *Octopus bimaculoides*, the SMPs of *Nautilus pompilius*, the EcMPs of *Argonauta argo*, the EcMPs of *Argonauta hians* indicated that the Argonauts do not use the same SMPs as EcMPs, while both SMPs and EcMPs were found to be present in the shell-less octopod *O. bimaculoides*.

Lastly, to identify conserved protein sequences, the annotated argonaut EcMPs were used as queries in reciprocal local BLASTp searches (blast-2.6.0-boost1.64_2), against four molluscan for which the shell matrix protein sequence data are already published as of June 2021 (71 *C. gigas* proteins (Zhang et al. 2012; Zhao et al. 2018); 159 *P. fucata* proteins (Takeuchi et al. 2016; Zhao et al. 2018); 311 *L. gigantea* proteins (Mann et al 2012)). Considering the even deeper divergence of the Conchiferans when compared to the Cephalopods, no hits were found. Therefore, we only showed and discussed the result of Search Setting 2 (Fig. 4).

**Fig. 4.**
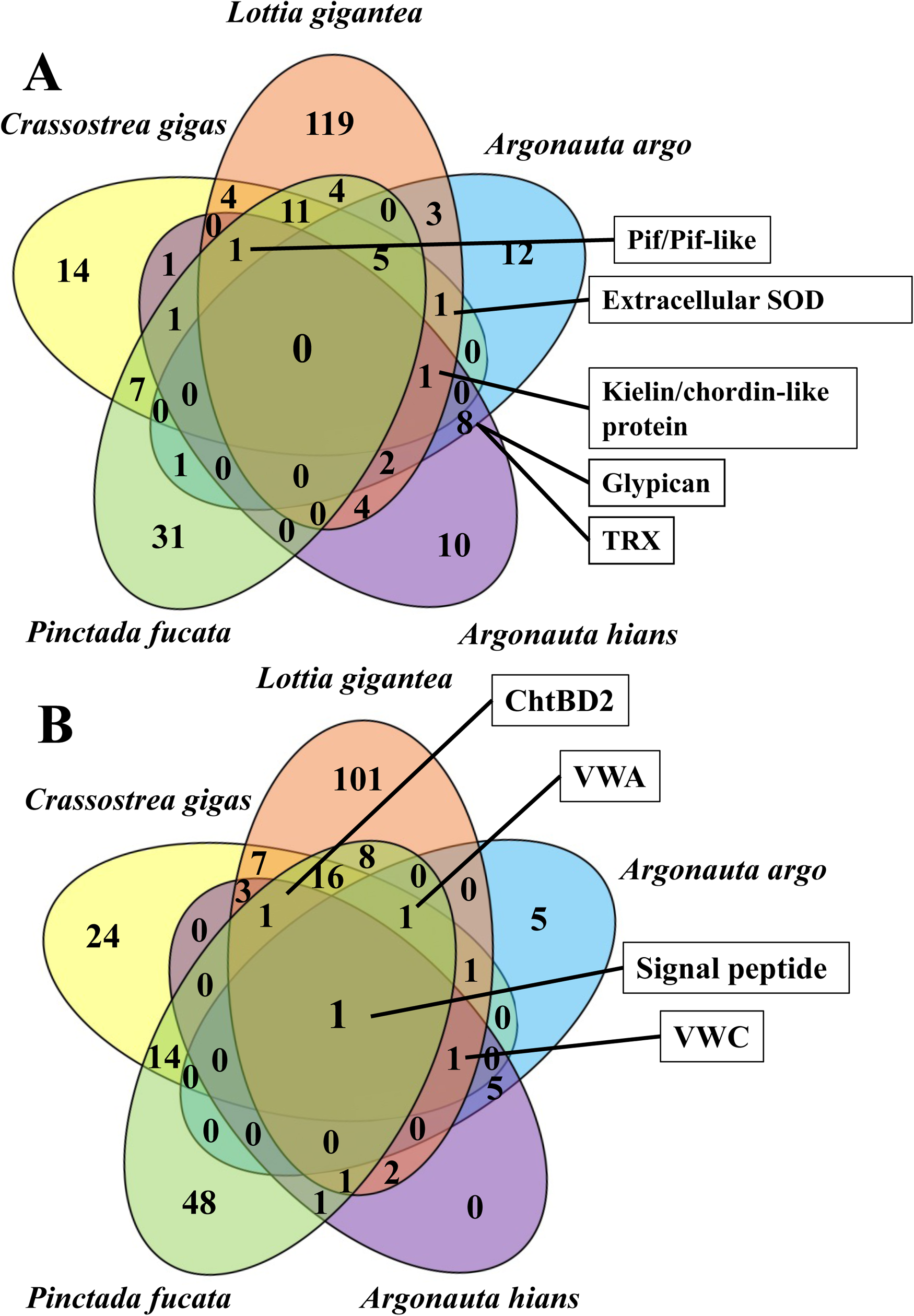
A diagram showing the result of the comparison of the SMPs and EcMPs among the Conchiferans. A. No shared proteins among the EcMPs of the Argonauts and the SMPs of the Conchiferans. B. Only the presence of the signal peptide domain is being shared among the various EcMPs of the Argonauta and SMPs of the Conchiferans.

### Phylogenetic analyses of the Shell Matrix Proteins

Phylogenetic analyses were conducted on a total of four EcMPs obtained in this study (PIF/BMSP-like protein, Superoxide dismutase (SOD), Thioredoxin (TRX), Glypican). To build single-gene trees based on orthologs, we performed webBLAST search using argonaut egg-case proteins. The sequences for the phylogenetic tree were collected covering the whole Lophotrochozoa clade. To perform multiple alignment of protein sequences, we utilized MAFFT v7.310 (Katoh et al. 2002), followed by the removal of ambiguously aligned sites using trimAl_v1.4beta (automated option) (Capella-Gutiérrez et al. 2009). Maximum likelihood phylogenetic inferences were executed on the software RAxMLGUI v2.0.5 (Silvestro et al. 2012; Stamatakis 2006) the rapid tree search setting with 1000 bootstrap replications under the best fit models (Glypican = LG + Γ + I; TRX = LG + Γ + I; Pif and Pif-like proteins = WAG + Γ; SOD = LG + Γ + I), which were tested using MEGA X (Kumar et al. 2018). Sequence data visualization and manipulation were done using the software MEGA X.

## Results

### 1. Acquisition of the EcMP-coding sequences of the two *Argonauta* species

We conducted PE100 transcriptome sequencing using the nine separated lanes (one lane per tissue sample), resulting in 24.9 to 28.9 million reads per run. After sequence assembly of reads from the six runs combined, 203,639 contigs were obtained for *A*. *argo*, with the largest contig being 23,502 bp-long, the average length of contigs 701.3 bp, and the N50 value of 1,094. Similarly, 164,654 contigs were obtained for *A. hians*, with the largest contig being 23,369 bp-long, the average length of contigs 781.5 bp, and the N50 value of 1,364.

Four runs of the LC-MS/MS mass spectrometer were conducted to analyze the two different fractions (hydrophilic and hydrophobic) of extracted proteins from the shell of *A. argo* and *A. hians*. A comparison between the obtained protein spectra from the MS/MS and the inferred protein spectra of the transcriptome contigs resulted in the identification of 109 proteins. Of these, 33 proteins were excluded from further analyses because they were identified as contaminants (Local BLASTn searches resulted in perfect matches to human, mouse, and bovine amino acid sequences). This gave us 21 soluble and 21 insoluble proteins for *A. argo*, and four soluble and 33 insoluble protein sequences for *A. hians*. Of these, 13 proteins in *A. argo* and three proteins in *A. hians* were found in both soluble and insoluble fractions, thus giving us the final result of 29 proteins in *A. argo* and 33 proteins in *A. hians*. Careful sequence analyses further identified that 16 (*A. argo*), 15 (*A. hians*) proteins had full length ORFs, and 16 (*A. argo*), 15 (*A. hians*) of them had signal peptides regardless of sequence completeness. Further domain searches identified the presence of domains in 29 of the 29 proteins of *A. argo*, and 31 of the 33 of *A. hians*. A complete list of the sequences obtained are given in Table 1 (for *A. argo*) and Table 2 (for *A. hians*). A schematic representation of the protein sequences with their domains of both species are shown in Fig. 2A (*A. argo*) and Fig. 2B (*A. hians*).

**Table 1.** The Eggcase Matrix Proteins in Argonauta argo.

| Accession | Soluble/Insoluble | Score (Soluble) | Score (Insoluble) | Mass | emPAI (Soluble) | emPAI (Insoluble) | BLAST top hit description | BLAST e-value (as of March 30, 2021) | SignalP |
| --- | --- | --- | --- | --- | --- | --- | --- | --- | --- |
| TRINITY_DN24597_c0_g1_i1_fr1 | Soluble/Insoluble | 257 | 1242 | 14360 | 1.17 | 2.64 | no hit |  | 5' incomplete |
| TRINITY_DN238_c0_g1_i1_fr5 | Soluble/Insoluble | 849 | 712 | 85299 | 0.71 | 0.64 | PREDICTED: glypican-6-like [*Octopus bimaculoides*]<br>Sequence ID: XP_014785616.1 | 0 | Minus |
| TRINITY_DN24867_c1_g1_i3_fr6 | Soluble/Insoluble | 454 | 2445 | 28243 | 0.71 | 2.8 | hypothetical protein OCBIM_22007538mg, partial<br>[*Octopus bimaculoides*]<br>Sequence ID: KOF68331.1 | 2.00E-52 | 5' incomplete |
| TRINITY_DN59231_c0_g1_i1_fr5 | Soluble | 94 |  | 10477 | 0.42 |  | PREDICTED: kielin/chordin-like protein [*Octopus bimaculoides*]<br>Sequence ID: XP_014785430.1 | 5.00E-24 | 5' incomplete |
| TRINITY_DN52918_c0_g1_i1_fr1 | Soluble/Insoluble | 108 | 82 | 31863 | 0.27 | 0.27 | PREDICTED: extracellular superoxide dismutase [Cu-Zn]-like [*Octopus bimaculoides*]<br>Sequence ID: XP_014772444.1 Length: 278 | 1.00E-65 | 5' incomplete |
| TRINITY_DN881_c6_g1_i1_fr2 | Soluble/Insoluble | 263 | 703 | 38362 | 0.22 | 0.64 | no hit |  | Plus |
| TRINITY_DN8120_c0_g1_i7_fr1 | Soluble/Insoluble | 406 | 5116 | 80938 | 0.21 | 1.13 | kielin/chordin-like protein [*Octopus sinensis*]<br>Sequence ID: XP_029634961.2 | 1.00E-173 | 5' incomplete |
| TRINITY_DN13749_c0_g1_i1_fr2 | Soluble | 79 |  | 27376 | 0.15 |  | PREDICTED: uncharacterized protein LOC106873894<br>[*Octopus bimaculoides*]<br>Sequence ID: XP_014776906.1 | 2.00E-29 | Plus |
| TRINITY_DN8120_c0_g2_i1_fr2 | Soluble/Insoluble | 67 | 128 | 28462 | 0.14 | 0.3 | PREDICTED: kielin/chordin-like protein [*Octopus bimaculoides*]<br>Sequence ID: XP_014785430.1 | 9.00E-39 | 5' incomplete |
| TRINITY_DN26366_c0_g1_i3_fr5 | Soluble/Insoluble | 47 | 103 | 43764 | 0.09 | 0.09 | PREDICTED: lysosomal aspartic protease-like isoform X1 [*Octopus bimaculoides*]<br>Sequence ID: XP_014771655.1 | 0 | 5' incomplete |
| TRINITY_DN1793_c1_g1_i1_fr5 | Soluble | 123 |  | 114938 | 0.07 |  | PREDICTED: protein disulfide-isomerase 2-like<br>[*Octopus bimaculoides*]<br>Sequence ID: XP_014785437.1 | 0 | Plus |
| TRINITY_DN951_c0_g1_i1_fr5 | Soluble/Insoluble | 60 | 78 | 59384 | 0.07 | 0.14 | lysosomal aspartic protease isoform X1 [*Octopus sinensis*]<br>Sequence ID: XP_029636147.1 | 0 | Minus |
| TRINITY_DN10130_c0_g1_i1_fr5 | Soluble/Insoluble | 48 | 59 | 59994 | 0.07 | 0.14 | ferritin-1, chloroplastic-like isoform X2 [*Octopus sinensis*]<br>Sequence ID: XP_029645066.1 | 7.00E-41 | Minus |
| TRINITY_DN726_c6_g1_i1_fr3 | Soluble | 70 |  | 67733 | 0.06 |  | probable peptidylglycine alpha-hydroxylating monooxygenase 1 [*Octopus sinensis*]<br>Sequence ID: XP_029646382.1 | 0 | Minus |
| TRINITY_DN1275_c0_g1_i10_fr5 | Soluble/Insoluble | 58 | 45 | 80924 | 0.05 | 0.05 | actin, cytoplasmic [*Octopus sinensis*]<br>Sequence ID: XP_029639042.1 | 0 | 5' incomplete |
| TRINITY_DN7032_c1_g1_i1_fr4 | Soluble | 57 |  | 71287 | 0.05 |  | collagen alpha-1(XII) chain [*Octopus sinensis*]<br>Sequence ID: XP_029655835.2 | 2.00E-130 | Plus |
| TRINITY_DN266_c2_g2_i2_fr2 | Soluble/Insoluble | 82 | 43 | 99137 | 0.04 | 0.04 | PREDICTED: 70 kDa neurofilament protein-like<br>[*Octopus bimaculoides*]<br>Sequence ID: XP_014776799.1 | 0 | 5' incomplete |
| TRINITY_DN432_c0_g1_i14_fr6 | Soluble/Insoluble | 68 | 32 | 114834 | 0.03 | 0.03 | kielin/chordin-like protein [*Octopus sinensis*]<br>Sequence ID: XP_036362851.1 | 2.00E-168 | Plus |
| TRINITY_DN4020_c0_g1_i1_fr1 | Soluble | 56 |  | 130420 | 0.03 |  | PREDICTED: lysosomal alpha-mannosidase-like<br>[*Octopus bimaculoides*]<br>Sequence ID: XP_014789463.1 | 0 | Minus |
| TRINITY_DN420_c8_g1_i1_fr5 | Soluble | 47 |  | 114467 | 0.03 |  | PREDICTED: glypican-3-like [*Octopus bimaculoides*]<br>Sequence ID: XP_014774973.1 | 0 | Minus |
| TRINITY_DN6962_c1_g1_i1_fr5 | Soluble | 65 |  | 179680 | 0.02 |  | PREDICTED: Golgi apparatus protein 1-like [*Octopus bimaculoides*]<br>Sequence ID: XP_014783789.1 | 0 | Minus |
| TRINITY_DN24867_c0_g1_i1_fr1 | Insoluble | 113 |  | 9681 |  | 3.5 | hypothetical protein OCBIM_22007538mg, partial<br>[*Octopus bimaculoides*]<br>Sequence ID: KOF68331.1 | 2.00E-52 | 5' incomplete |
| TRINITY_DN33718_c0_g1_i1_fr2 | Insoluble | 521 |  | 8745 |  | 1.28 | PREDICTED: kielin/chordin-like protein [*Octopus bimaculoides*]<br>Sequence ID: XP_014789136.1 | 1.00E-21 | 5' incomplete |
| TRINITY_DN13749_c0_g1_i2_fr3 | Insoluble | 96 |  | 33005 |  | 0.41 | PREDICTED: uncharacterized protein LOC106873894<br>[*Octopus bimaculoides*]<br>Sequence ID: XP_014776906.1 | 1.00E-28 | Minus |
| TRINITY_DN15083_c0_g2_i1_fr5 | Insoluble | 54 |  | 22083 |  | 0.4 | histone H4-like protein [*Labeo rohita*]<br>Sequence ID: RXN31026.1 | 3.00E-66 | 5' incomplete |
| TRINITY_DN25002_c0_g1_i4_fr6 | Insoluble | 67 |  | 21890 |  | 0.19 | PREDICTED: von Willebrand factor C and EGF domain-containing protein-like, partial [*Octopus bimaculoides*]<br>Sequence ID: XP_014788139.1 | 1.00E-58 | 5' incomplete |
| TRINITY_DN4158_c0_g1_i1_fr2 | Insoluble | 64 |  | 40193 |  | 0.1 | PREDICTED: uncharacterized protein LOC106872312<br>[*Octopus bimaculoides*]<br>Sequence ID: XP_014774739.1 | 2.00E-25 | Plus |
| TRINITY_DN34538_c0_g1_i1_fr6 | Insoluble | 52 |  | 46573 |  | 0.08 | no hit |  | Plus |
| TRINITY_DN9416_c1_g1_i10_fr2 | Insoluble | 76 |  | 98614 |  | 0.04 | zinc finger protein 271-like [*Octopus sinensis*]<br>Sequence ID: XP_036364921.1 | 1.00E-162 | Plus |

**Table 2.** The Eggcase Matrix Proteins in Argonauta hians.

| Accession | Soluble/Insoluble | Score (Soluble) | Score (Insoluble) | Mass | emPAI (Soluble) | emPAI (Insoluble) | BLAST top hit description | BLAST e-value (as of March 30, 2021) | SignalP |
| --- | --- | --- | --- | --- | --- | --- | --- | --- | --- |
| TRINITY_DN153_c0_g1_i4_fr4 | Soluble |  | 106 | 23491 |  | 0.17 | PREDICTED: kielin/chordin-like protein [*Octopus bimaculoides*]<br>Sequence ID: XP_014789136.1 | 4.00E-49 | 5' incomplete |
| TRINITY_DN568_c0_g1_i14_fr3 | Soluble/Insoluble | 69 | 438 | 25199 | 0.16 | 0.56 | PREDICTED: kielin/chordin-like protein [*Octopus bimaculoides*]<br>Sequence ID: XP_014785430.1 | 4.00E-26 | 5' incomplete |
| TRINITY_DN2436_c0_g1_i1_fr5 | Soluble/Insoluble | 74 | 575 | 40252 | 0.1 | 0.6 | no hit |  | Plus |
| TRINITY_DN367_c0_g1_i1_fr3 | Soluble/Insoluble | 81 | 37 | 188560 | 0.02 | 0.02 | PREDICTED: protein disulfide-isomerase 2-like [*Octopus bimaculoides*]<br>Sequence ID: XP_014785437.1 | 0 | Plus |
| TRINITY_DN568_c0_g1_i10_fr1 | Insoluble |  | 967 | 24384 |  | 0.85 | PREDICTED: kielin/chordin-like protein, partial [*Octopus bimaculoides*]<br>Sequence ID: XP_014786769.1 | 4.00E-24 | 5' incomplete |
| TRINITY_DN153_c0_g1_i1_fr6 | Insoluble |  | 2756 | 25358 |  | 0.81 | hypothetical protein OCBIM_22012348mg, partial [*Octopus bimaculoides*]<br>Sequence ID: KOF66424.1 | 2.00E-29 | 5' incomplete |
| TRINITY_DN3795_c0_g1_i1_fr2 | Insoluble |  | 54 | 8008 |  | 0.57 | PREDICTED: kielin/chordin-like protein, partial [*Octopus bimaculoides*]<br>Sequence ID: XP_014786769.1 | 1.00E-12 | 5' incomplete |
| TRINITY_DN22369_c0_g1_i1_fr6 | Insoluble |  | 86 | 34360 |  | 0.55 | histone H4-like [*Carassius auratus*]<br>Sequence ID: XP_026072634.1 | 3.00E-66 | Minus |
| TRINITY_DN39376_c0_g1_i2_fr2 | Insoluble |  | 64 | 17819 |  | 0.52 | keratin, type I cytoskeletal 13-like [*Seriola dumerili*]<br>Sequence ID: XP_022607894.1 | 8.00E-98 | 5' incomplete |
| TRINITY_DN153_c1_g1_i10_fr6 | Insoluble |  | 77 | 18900 |  | 0.49 | mucin-5B-like, partial [*Octopus sinensis*]<br>Sequence ID: XP_036362850.1 | 1.00E-35 | 5' incomplete |
| TRINITY_DN4045_c3_g2_i3_fr6 | Insoluble |  | 32 | 9149 |  | 0.49 | no hit |  | 5' incomplete |
| TRINITY_DN121_c3_g1_i2_fr1 | Insoluble |  | 85 | 9170 |  | 0.48 | no hit |  | 5' incomplete |
| TRINITY_DN5301_c0_g1_i5_fr1 | Insoluble |  | 155 | 19305 |  | 0.47 | PREDICTED: von Willebrand factor C and EGF domain-containing protein-like, partial [*Octopus bimaculoides*]<br>Sequence ID: XP_014788139.1 | 3.00E-52 | 5' incomplete |
| TRINITY_DN153_c1_g2_i1_fr4 | Insoluble |  | 100 | 10497 |  | 0.41 | mucin-5B-like, partial [*Octopus sinensis*]<br>Sequence ID: XP_036362850.1 | 1.00E-35 | 5' incomplete |
| TRINITY_DN18513_c0_g1_i2_fr2 | Insoluble |  | 430 | 13018 |  | 0.33 | no hit (serine-rich) |  | 5' incomplete |
| TRINITY_DN616_c0_g1_i13_fr6 | Insoluble |  | 137 | 44733 |  | 0.29 | kielin/chordin-like protein [*Octopus sinensis*]<br>Sequence ID: XP_036358531.1 | 9.00E-97 | 5' incomplete |
| TRINITY_DN212_c4_g1_i1_fr1 | Insoluble |  | 206 | 82200 |  | 0.26 | PREDICTED: glypican-6-like [*Octopus bimaculoides*]<br>Sequence ID: XP_014785616.1 | 0 | Minus |
| TRINITY_DN6154_c0_g1_i13_fr2 | Insoluble |  | 118 | 16951 |  | 0.24 | mucin-5B-like, partial [*Octopus sinensis*]<br>Sequence ID: XP_036362850.1 | 2.00E-22 | 5' incomplete |
| TRINITY_DN153_c1_g1_i3_fr5 | Insoluble |  | 271 | 19397 |  | 0.21 | PREDICTED: kielin/chordin-like protein [*Octopus bimaculoides*]<br>Sequence ID: XP_014789136.1 | 4.00E-49 | 5' incomplete |
| TRINITY_DN902_c2_g1_i1_fr2 | Insoluble |  | 44 | 20142 |  | 0.2 | no hit |  |  |
| TRINITY_DN1775_c0_g1_i1_fr1 | Insoluble |  | 48 | 21317 |  | 0.19 | PREDICTED: histone H2A-like [*Fopius arisanus*]<br>Sequence ID: XP_011311067.1 | 5.00E-78 | Minus |
| TRINITY_DN17_c1_g3_i1_fr4 | Insoluble |  | 53 | 47406 |  | 0.17 | PREDICTED: uncharacterized protein LOC106872312 [*Octopus bimaculoides*]<br>Sequence ID: XP_014774739.1 | 6.00E-24 | Minus |
| TRINITY_DN767_c1_g1_i3_fr4 | Insoluble |  | 344 | 25389 |  | 0.16 | PREDICTED: kielin/chordin-like protein [*Octopus bimaculoides*]<br>Sequence ID: XP_014790947.1 | 5.00E-71 | 5' incomplete |
| TRINITY_DN337_c1_g1_i10_fr4 | Insoluble |  | 48 | 37578 |  | 0.11 | PREDICTED: uncharacterized protein LOC106872312 [*Octopus bimaculoides*]<br>Sequence ID: XP_014774739.1 | 5.00E-24 | Plus |
| TRINITY_DN1327_c0_g2_i1_fr4 | Insoluble |  | 64 | 88359 |  | 0.09 | PREDICTED: actin, adductor muscle [*Octopus bimaculoides*]<br>Sequence ID: XP_014771927.1 | 0 | Minus |
| TRINITY_DN3465_c0_g3_i1_fr4 | Insoluble |  | 43 | 42856 |  | 0.09 | soma ferritin-like [*Octopus sinensis*]<br>Sequence ID: XP_029644988.1 | 3.00E-44 | Plus |
| TRINITY_DN17695_c0_g2_i1_fr1 | Insoluble |  | 56 | 62738 |  | 0.06 | PREDICTED: uncharacterized protein LOC106873894 [*Octopus bimaculoides*]<br>Sequence ID: XP_014776906.1 | 2.00E-24 | Minus |
| TRINITY_DN1213_c0_g1_i1_fr3 | Insoluble |  | 53 | 69122 |  | 0.06 | PREDICTED: elongation factor 1-alpha-like [*Octopus bimaculoides*]<br>Sequence ID: XP_014782958.1 | 0 | Minus |
| TRINITY_DN358_c0_g1_i1_fr6 | Insoluble |  | 30 | 63664 |  | 0.06 | PREDICTED: lysosomal aspartic protease-like isoform X1 [*Octopus bimaculoides*]<br>Sequence ID: XP_014771655.1 | 0 | Plus |
| TRINITY_DN2622_c1_g4_i4_fr4 | Insoluble |  | 56 | 75220 |  | 0.05 | PREDICTED: protein PIF-like [*Octopus bimaculoides*]<br>Sequence ID: XP_014787620.1 | 0 | Plus |
| TRINITY_DN1286_c0_g1_i4_fr5 | Insoluble |  | 49 | 87538 |  | 0.04 | PREDICTED: von Willebrand factor C and EGF domain-containing protein-like, partial [*Octopus bimaculoides*]<br>Sequence ID: XP_014788139.1 | 1.00E-149 | Plus |
| TRINITY_DN3214_c0_g1_i1_fr2 | Insoluble |  | 111 | 145903 |  | 0.03 | hypothetical protein T12_2372 [*Trichinella patagoniensis*]<br>Sequence ID: KRY08652.1 | 1.00E-18 | Minus |
| TRINITY_DN3475_c0_g3_i2_fr2 | Insoluble |  | 42 | 141608 |  | 0.03 | PREDICTED: pantetheinase-like [*Octopus bimaculoides*]<br>Sequence ID: XP_014777355.1 | 0 | Plus |

### 2. Identification of shared proteins between the shell of Conchiferans and the eggcase of the Argonauts

In order to understand which proteins are shared in the eggcase matrix of the two species of *Argonauta*, we conducted reciprocal local BLASTp searches between the newly identified EcMPs sequences obtained from the two species (29 proteins for *A. argo* and 33 for *A. hians*). Because of the possible recent divergence of the two argonaut species, we conducted BLASTp searches under a stringent setting (Search Setting 1; e-value <1e-50 and threshold ≥40%) in order to avoid false positives in homology searches. Through the stringent searches, 15 proteins were found to be present only in *A. argo* (Fig. 2A), and 16 in *A. hians* (Fig. 2B), while 14 proteins of *A. argo* were found as probable homologs of 17 proteins of *A. hians* (Fig. 2C). The list of proteins, categorized as those shared between the two species and species-specific ones, are shown in Table 1. Meanwhile, for confirmation, searches under the less stringent Search Setting 2 (e-value <1e-5) were also carried out. These searches gave us eight proteins specific to *A. argo*, and 11 to *A. hians*, and 21 proteins of *A. argo* shared with 22 proteins of *A. hians* (Supplementary Fig. 1).

Next, we carried out similar comparative analyses using local BLASTp among the EcMPs of the two argonauts used in this study and the gene models obtained from the genome of the shell-less Octopodiform *Octopus bimaculoides* (the octopus). From the local BLASTp searches under Search Setting 2, 25 proteins of the EcMPs in *A. argo* and 27 proteins of the EcMPs in *A. hians* were found to be putatively homologous to 669 and 143 proteins into the octopus genome, respectively (Fig. 3). When the zinc finger-domain containing protein, C2H2 Zinc-Finger protein, found as one of the EcMPs of *A. argo* (not found in the EcMPs of *A. hians*) was excluded from the search, the putative octopus gene homologs of *A. argo* genes found in the octopus genome became 199 (Fig. 3; Supplementary Fig. 2). Adding the SMPs data of the Nautiliid *Nautilus pompilius* (the nautilus) (Setiamarga et al. 2020) to the analyses above also gave us an interesting result (Fig. 3). First, no SMP was found to be homologous among the three species, while one (Pif-like/Laminin G3) was found to be shared between SMPs of the nautilus and EcMPs of *A. hians*. Meanwhile, putative homologs of 15 of the 47 SMP-coding genes of the nautilus (Setiamarga et al. 2020) were found in the octopus genome, including Pif-like/Laminin G3 found in the EcMPs of *A. hians*. It is to be noted that, while we could not find the said protein in the EcMPs of *A. argo*, we did find similar proteins in the mantle transcriptome data of *A. argo*.

In order to find out if there was any conserved protein and protein domain among the SMPs of the Conchiferans and the EcMPs of the argonauts, reciprocal local BLASTp searches on the argonaut EcMPs and the SMP data of selected three Conchiferans for which exhaustive SMP data are available (the pacific oyster *Crassostrea gigas*, the pearl oyster *Pinctada fucata*, and the limpet *Lottia gigantea*) were conducted with the threshold of 1≤e-5 (Search Setting 2) (Fig. 4).

From the searches, we found no proteins conserved among the SMPs of the Conchiferans and the EcMPs of the Argonauts. However, we also found several proteins shared among some of the Conchiferans and one or both species of the Argonauts. For example, a full-length ORF with a signal peptide of the Pif-like protein (Laminin G3) was found in *A. hians*. The protein consists of three consecutive chitin-binding domains (SMART ChtBD2 domain, SM000494) followed by a Laminin_G3 domain (PF13385). Although the ORF was only detected in the EcMPs of *A. hians*, a contig (DN73320_c0_g1_i1_fr4) matching the first half of the 123 amino acids (99%) and a contig (DN41781_c0_g1_i1_fr1) matching the second half of the 82 amino acids (98%) were obtained from the transcriptome of *A. argo*. Although in this study we only found homologous sequences with a similar domain structure in *Pinctada fucata* (Matsuura et al. 2018), the presence of the Pif-like/Laminin G3 protein in the shell matrix were also reported from the nautilus (Setiamarga et al. 2020), a pectinid bivalve *Argopecten purpuratus* (Li et al. 2018), and the pond snail *Lymnaea stagnalis* (Ishikawa et al. 2020).

Two enzymes known to be involved in antioxidizing processes, Thioreductase (TRX) and Superoxide Dismutase (SOD), were also detected in both the shell matrix of the Conchiferans, and the EcMPs of the Argonauts. TRX was found in both *A. argo* and *A. hians*. Three thioredoxin domains following a signal peptide cover the entire length of the protein in DN1793_c1_g1_i1_fr5 of *A. argo*, and DN367_c0_g1_i1_fr3 in *A. hians*. Although in our analysis, we only found TRX in the two Argonauts, previously, Marie et al. (2017) reported its presence in the SMPs of a freshwater mussel, *Villosa lienosa*. A full length of SOD sequence was found in *A. argo* (DN52918_c0_g1_i1_fr1), and its putative homologs were found in the SMP data of *Lottia gigantea* and *Crassostrea gigas*. The protein Ferritin, which was the principal protein for iron storage, was also found to be shared among the two Argonaut species (*A. argo*: partial, DN10130_c0_g1_i1_fr5; *A. hians*: full length, DN3465_c0_g3_i1_fr4) and *P. fucata*. The involvement of Ferritin in shell formation has been suggested by various previous studies (e.g., *Pinctada fucata:* Zhang *et al*. (2003); *Haliotis discus*: De Zoysa and Lee 2007).

Peptides translated from standard housekeeping genes such as *actin* (both argonauts), *histone* H2 (only *A. hians*), *histone* H4 (in both argonauts), and *neurofilament* (only in *A. argo*) were also detected in the eggcase matrix. This observation supports what was reported previously by Oudot et al. (2020b). The presence of housekeeping proteins in the shell matrix is not surprising, although it is intriguing because no study has addressed what they are doing there. Housekeeping proteins have also been reported to be present in the shell matrix of *Crassostrea angulata* and *C. hongkongensis* (Upadhyay et al. 2016), *Euhadra quaesita* (Shimizu et al. 2019), and *Spirula spirula* (Oudot et al. 2020a).

Based on the order of emPAI frequency, the most abundantly present protein was kielin/chordin-like proteins. These are von Willebrand factor type C (VWC)-containing proteins that structurally only have either single or multiple VWC domains (Table 1, Fig. 2). Nine contigs of this protein were found in *A. argo*, while 13 was found in *A. hians*. Kielin/chordin-like proteins were also found in *L. gigantea* and *C. gigas*. The next most frequently observed protein was Glypican, which is a member of the glypican-related integral membrane proteoglycan family (GRIPS). The GRIPS proteins contain a core protein anchored to the cytoplasmic membrane via a glycosyl phosphatidylinositol (GPI) linkage. Two different protein contigs were found in *A. argo* (DN420_c8_g1_i1_fr5 and DN238_c0_g1_i1_fr5). The first contig contains a probable GPI site on the N-terminal side of the Glypican domain (Fig. 2A). In *A. hians*, only one Glypican protein contig (DN212_c4_g1_i1_fr1) was found (Fig. 2B).

### 3. Gene expression patterns of EcMP-coding genes based on the transcriptome data

The Pif-like protein detected as one of the EcMPs, LamG3/Pif-like, was very highly expressed in the epithelial tissues of *A. argo* (kallisto abundance values 1020.02, 1201.66, and 832.5 in the first, second arms, and mantle, respectively). Meanwhile, the TRX was found to be biased in the epithelial tissues (kallisto abundances > 28 in the arms and mantle, but 4∼1.8 in the other tissues). The SOD was only found in the first arm and mantle (kallisto abundances = 1.44 and 1.03). Glypican was expressed throughout all tissues (kallisto abundances 8.6∼24.2). The summary of the place of expression of the 29 EcMP-coding genes of *A. argo* based on transcriptome data is presented in Table 3.

**Table 3.** Transcriptome expression abundance in *Argonauta argo*.

| Accession | BLAST top hit description | Detail | abundance<br>1st Arm | abundance<br>2nd Arm | abundance<br>Eye | abundance<br>Gill Heart | abundance<br>Heart | abundance<br>Mantle |
| --- | --- | --- | --- | --- | --- | --- | --- | --- |
| TRINITY_DN24597_c0_g1_i1 | no hit |  | 0.282806 | 0.487961 | 0 | 0 | 0 | 0.530571 |
| TRINITY_DN238_c0_g1_i1 | PREDICTED: glypican-6-like [Octopus bimaculoides], Sequence ID: XP_014785616.1 | Glypican | 16.1295 | 24.2499 | 1.37913 | 15.0179 | 7.35824 | 8.6354 |
| TRINITY_DN24867_c1_g1_i3 | hypothetical protein OCBIM_22007538mg, partial [Octopus bimaculoides], Sequence ID: KOF68331.1 |  | 0.754866 | 1.21665 | 0 | 0.0707456 | 0 | 0.544948 |
| TRINITY_DN24867_c1_g1_i3 | PREDICTED: kielin/chordin-like protein [Octopus bimaculoides], Sequence ID: XP_014785430.1 |  | 0.839783 | 1.15131 | 0 | 0 | 0 | 0.00E+00 |
| TRINITY_DN52918_c0_g1_i1 | PREDICTED: extracellular superoxide dismutase [Cu-Zn]-like [Octopus bimaculoides], Sequence ID: XP_014772444.1Length: 278 | SOD | 1.44415 | 0.227733 | 0 | 0.113231 | 0 | 1.03E+00 |
| TRINITY_DN881_c6_g1_i1 | no hit |  | 210.072 | 54.4012 | 75.9833 | 527.697 | 686.373 | 4.41E+01 |
| TRINITY_DN8120_c0_g1_i7 | kielin/chordin-like protein [Octopus sinensis], Sequence ID: XP_029634961.2 |  | 1.30374 | 2.45628 | 0 | 0 | 0 | 1.17E+00 |
| TRINITY_DN13749_c0_g1_i1 | PREDICTED: uncharacterized protein LOC106873894 [Octopus bimaculoides], Sequence ID: XP_014776906.1 |  | 6.02102 | 1.06272 | 0.725911 | 0.551878 | 1.29541 | 6.37E-01 |
| TRINITY_DN8120_c0_g2_i1 | PREDICTED: kielin/chordin-like protein [Octopus bimaculoides], Sequence ID: XP_014785430.1 |  | 0 | 1.49045 | 0 | 0 | 0 | 5.58E-01 |
| TRINITY_DN26366_c0_g1_i3 | PREDICTED: lysosomal aspartic protease-like isoform X1 [Octopus bimaculoides], Sequence ID: XP_014771655.1 |  | 0.657002 | 0.61553 | 0.125311 | 0 | 0 | 2.07443 |
| TRINITY_DN1793_c1_g1_i1 | PREDICTED: protein disulfide-isomerase 2-like [Octopus bimaculoides], Sequence ID: XP_014785437.1 | TRX | 30.8582 | 75.5665 | 3.34925 | 3.98771 | 1.82726 | 2.87E+01 |
| TRINITY_DN951_c0_g1_i1 | lysosomal aspartic protease isoform X1 [Octopus sinensis], Sequence ID: XP_029636147.1 |  | 57.7914 | 83.5223 | 127.047 | 594.462 | 582.984 | 5.20E+01 |
| TRINITY_DN10130_c0_g1_i1 | ferritin-1, chloroplastic-like isoform X2 [Octopus sinensis], Sequence ID: XP_029645066.1 |  | 2.57981 | 6.86812 | 3.787 | 1.79804 | 0.121306 | 2.47E+00 |
| TRINITY_DN726_c6_g1_i1 | probable peptidylglycine alpha-hydroxylating monooxygenase 1 [Octopus sinensis], Sequence ID: XP_029646382.1 |  | 239.5 | 66.9307 | 18.4668 | 109.174 | 39.0137 | 2.41E+01 |
| TRINITY_DN1275_c0_g1_i10 | actin, cytoplasmic [Octopus sinensis], Sequence ID: XP_029639042.1 |  | 38.2882 | 137.047 | 293.293 | 294.583 | 45.0621 | 1.05E+01 |
| TRINITY_DN7032_c1_g1_i1 | collagen alpha-1(XII) chain [Octopus sinensis], Sequence ID: XP_029655835.2 |  | 391.494 | 122.157 | 0 | 10.5932 | 4.63763 | 1.77E+03 |
| TRINITY_DN266_c2_g2_i2 | PREDICTED: 70 kDa neurofilament protein-like [Octopus bimaculoides], Sequence ID: XP_014776799.1 |  | 198.831 | 195.846 | 0 | 0 | 0 | 463.447 |
| TRINITY_DN432_c0_g1_i14 | kielin/chordin-like protein [Octopus sinensis], Sequence ID: XP_036362851.1 |  | 0.00720252 | 127.413 | 0 | 0 | 0 | 0.093546 |
| TRINITY_DN4020_c0_g1_i1 | PREDICTED: lysosomal alpha-mannosidase-like [Octopus bimaculoides], Sequence ID: XP_014789463.1 |  | 24.4716 | 16.228 | 1.01886 | 23.3563 | 8.92014 | 27.2065 |
| TRINITY_DN420_c8_g1_i1 | PREDICTED: glypican-3-like [Octopus bimaculoides], Sequence ID: XP_014774973.1 |  | 14.773 | 20.943 | 6.57756 | 12.1627 | 6.92061 | 17.5107 |
| TRINITY_DN6962_c1_g1_i1 | PREDICTED: Golgi apparatus protein 1-like [Octopus bimaculoides], Sequence ID: XP_014783789.1 |  | 7.63314 | 11.7315 | 1.97418 | 5.52828 | 2.56404 | 5.52923 |
| TRINITY_DN24867_c0_g1_i1 | hypothetical protein OCBIM_22007538mg, partial [Octopus bimaculoides], Sequence ID: KOF68331.1 |  | 0.743906 | 0 | 0 | 0 | 0 | 2.8435 |
| TRINITY_DN33718_c0_g1_i1 | PREDICTED: kielin/chordin-like protein [Octopus bimaculoides], Sequence ID: XP_014789136.1 |  | 2.5172 | 2.30806 | 0 | 0 | 0 | 0.00E+00 |
| TRINITY_DN13749_c0_g1_i2 | PREDICTED: uncharacterized protein LOC106873894 [Octopus bimaculoides], Sequence ID: XP_014776906.1 |  | 2.47131 | 0 | 0 | 1.08692 | 0.271559 | 1.95E-01 |
| TRINITY_DN15083_c0_g2_i1 | histone H4-like protein [Labeo rohita], Sequence ID: RXN31026.1 |  | 20.596 | 20.2651 | 0.922869 | 33.7763 | 26.0391 | 14.6815 |
| TRINITY_DN25002_c0_g1_i4 | PREDICTED: von Willebrand factor C and EGF domain-containing protein-like, partial [Octopus bimaculoides], Sequence ID: XP_014788139.1 |  | 0.627458 | 0.350163 | 0.0941585 | 0 | 0 | 0.504363 |
| TRINITY_DN4158_c0_g1_i1 | PREDICTED: uncharacterized protein LOC106872312 [Octopus bimaculoides], Sequence ID: XP_014774739.1 |  | 145.599 | 65.8228 | 0 | 2.3723 | 0.410338 | 312.937 |
| TRINITY_DN34538_c0_g1_i1 | no hit |  | 0.297561 | 0.539167 | 1.4044 | 0 | 0 | 0.164701 |
| TRINITY_DN9416_c1_g1_i10 | zinc finger protein 271-like [Octopus sinensis], Sequence ID: XP_036364921.1 |  | 2.61831E-06 | 0 | 0.110231 | 0 | 0 | 1.51E+00 |

When we checked if there was any highly expressed protein for shell pigmentation or calcification commonly found as Conchiferan SMP, we found tyrosinases with specific expressions in the epithelia of *A. argo* (Fig. 6). When we checked the expression patterns of the contigs of Tyrosinase, we found that contig DN84_c0_g1 in *A. argo* showed a very high expression only in arms (TPM>1,000), but zero in eyes, heart and gill heart. The ortholog candidate sequence found in the transcriptome data of *A. hians*, DN16_c0_g1, was also highly expressed in the arms (Fig. 6). We also found that another contig DN4172_c0_g1 was specifically expressed in epithelial tissues of *A. hians* (Fig. 6).

### 4. Molecular phylogenetics of some of the Argonaut EcMPs

In order to look into the molecular evolutionary history of EcMPs, we carried out maximum likelihood phylogenetic inferences of four proteins found in the eggcase of the two species of *Argonauta* targeted in this study, (Laminin G3/Pif-like protein, Thioredoxin, Superoxide Dismutase, and Glypican). We also analyzed Tyrosinase, an enzyme known as a major SMP in Conchiferans, although the enzyme was not detected in the eggcase matrix proteome data of the Argonauts. For the analyses, homologs of these proteins were obtained from GenBank, based on various previous studies.

Five sequences of LamininG3/Pif-like protein were found in both species of Argonauts analyzed in this study, of which three sequences were of full length (Fig. 5A). While Pif proteins identified as a Conchiferan SMP usually have two domains, (von Willebrand factor type A (VWA) and chitin-binding (ChtBd)), the argonaut Pif-like protein had only two domains, the ChtBd and laminin G3 (LamG3) domains, even in full-length proteins.The result of our phylogenetic analysis placed the five sequences together with other similarly structured Laminin G3/Pif-like proteins of the Conchiferans, forming a monophyletic group with the sequences of the gastropods and bivalves, besides other cephalopods such as the cuttlefish and even the shell-less octopus (Fig. 5C). Laminin G3/ Pif-like protein forms a monophyletic clade, separated from the other two monophyletic clades of major Conchiferan SMPs, the Pif and BMSP proteins, which have different domains and domain arrangements from each other (Fig. 5C). The domain structures of Laminin G3/Pif-like, Pif, and BMSP are shown in Fig. 5A.

**Fig. 5.**
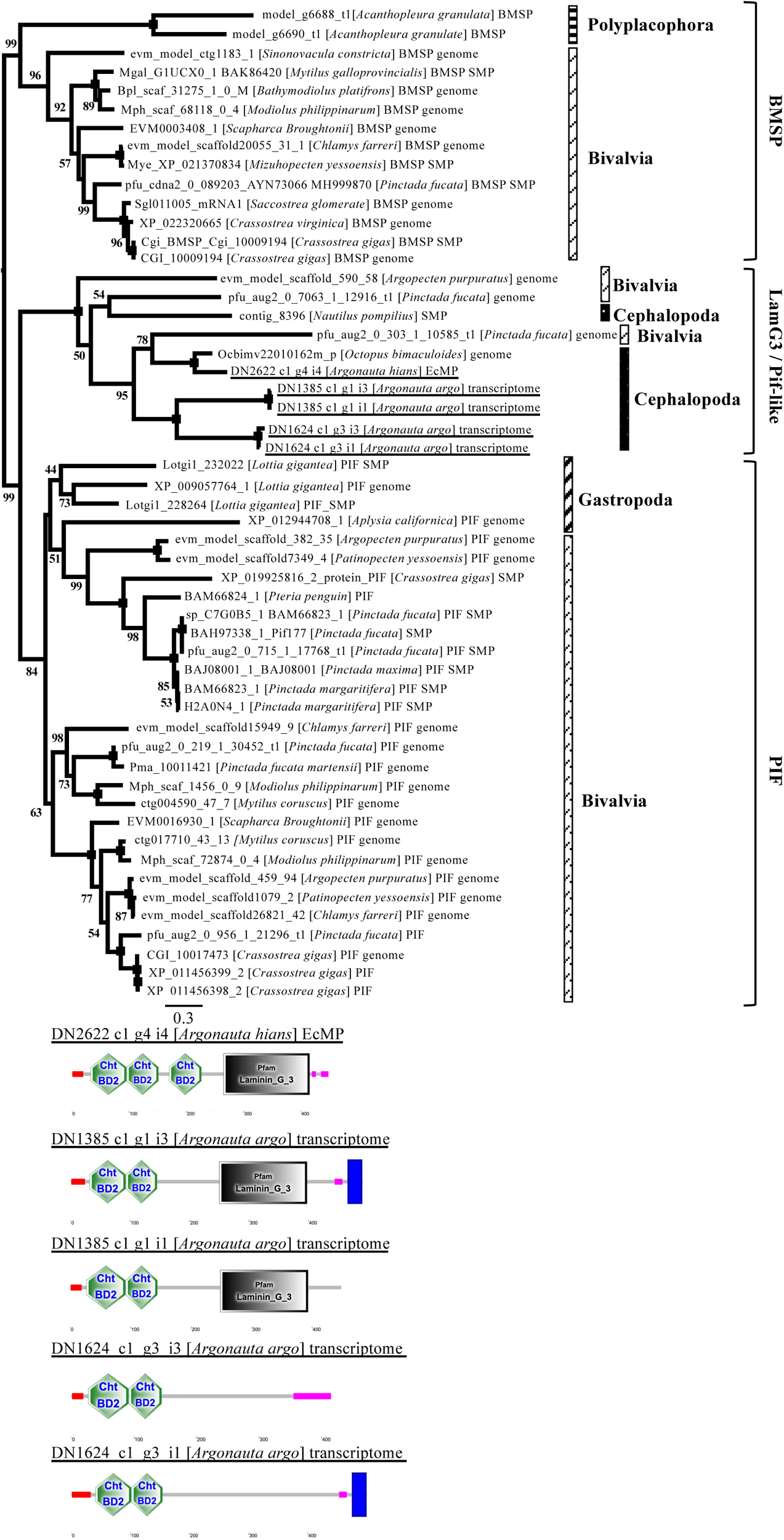
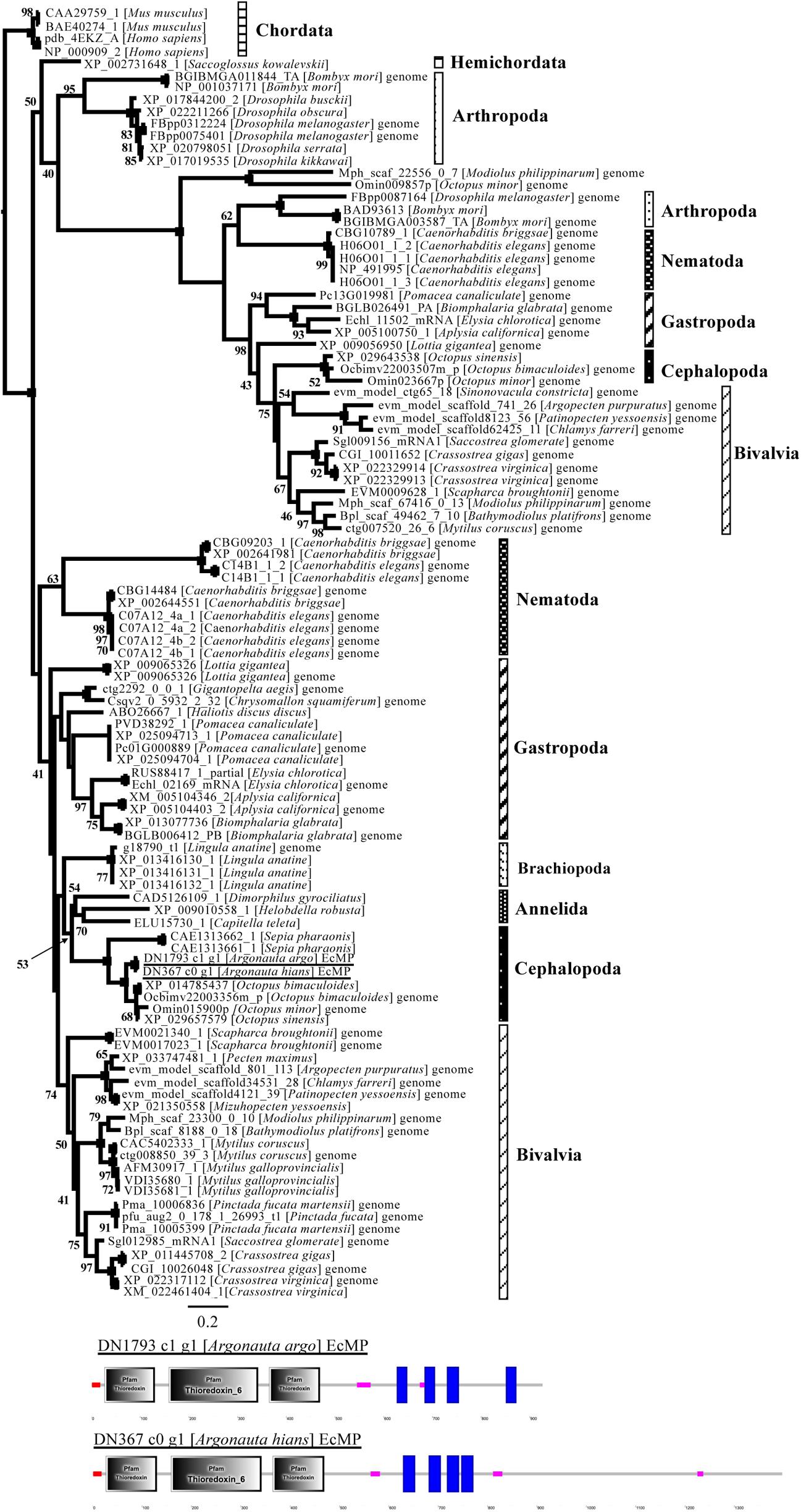
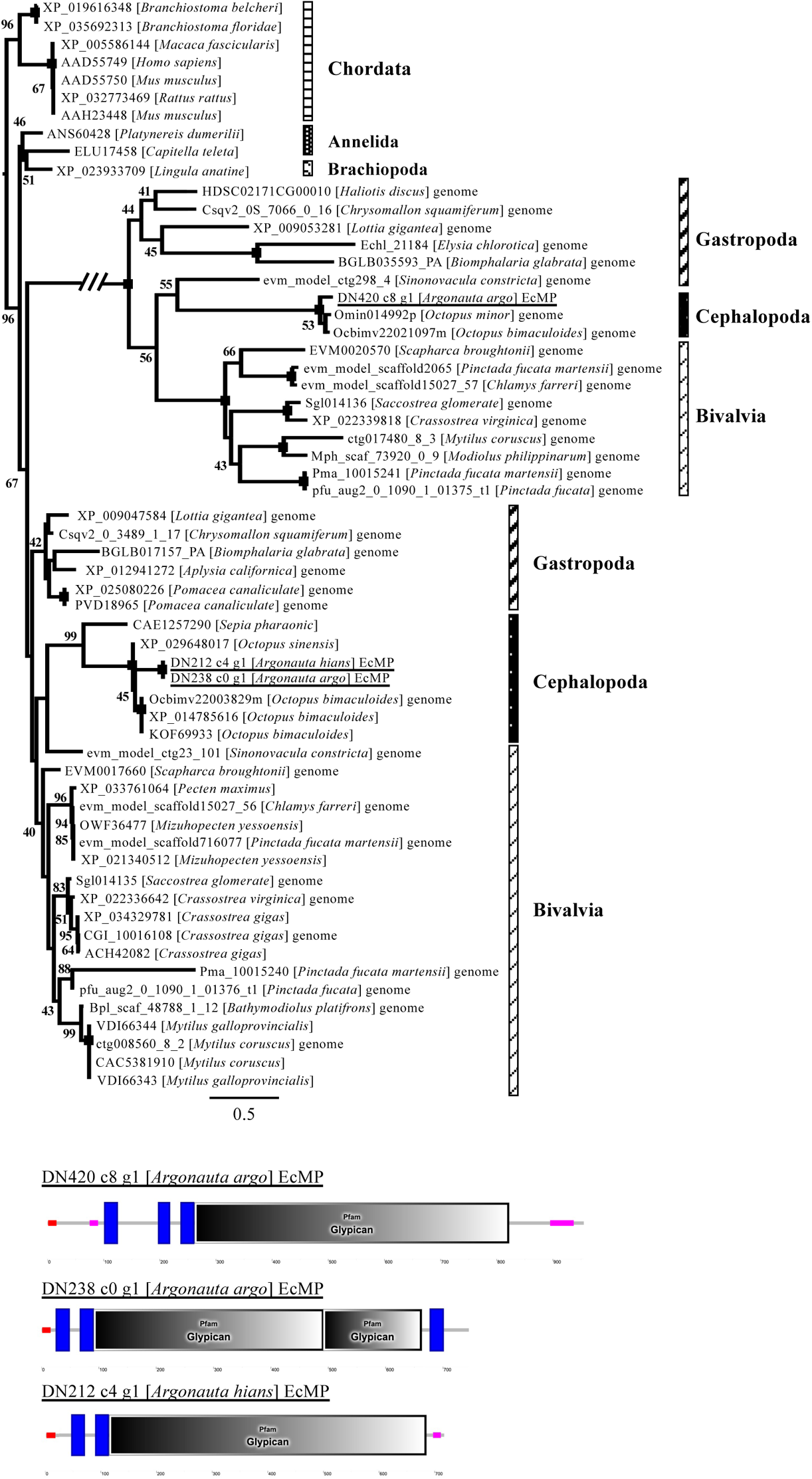
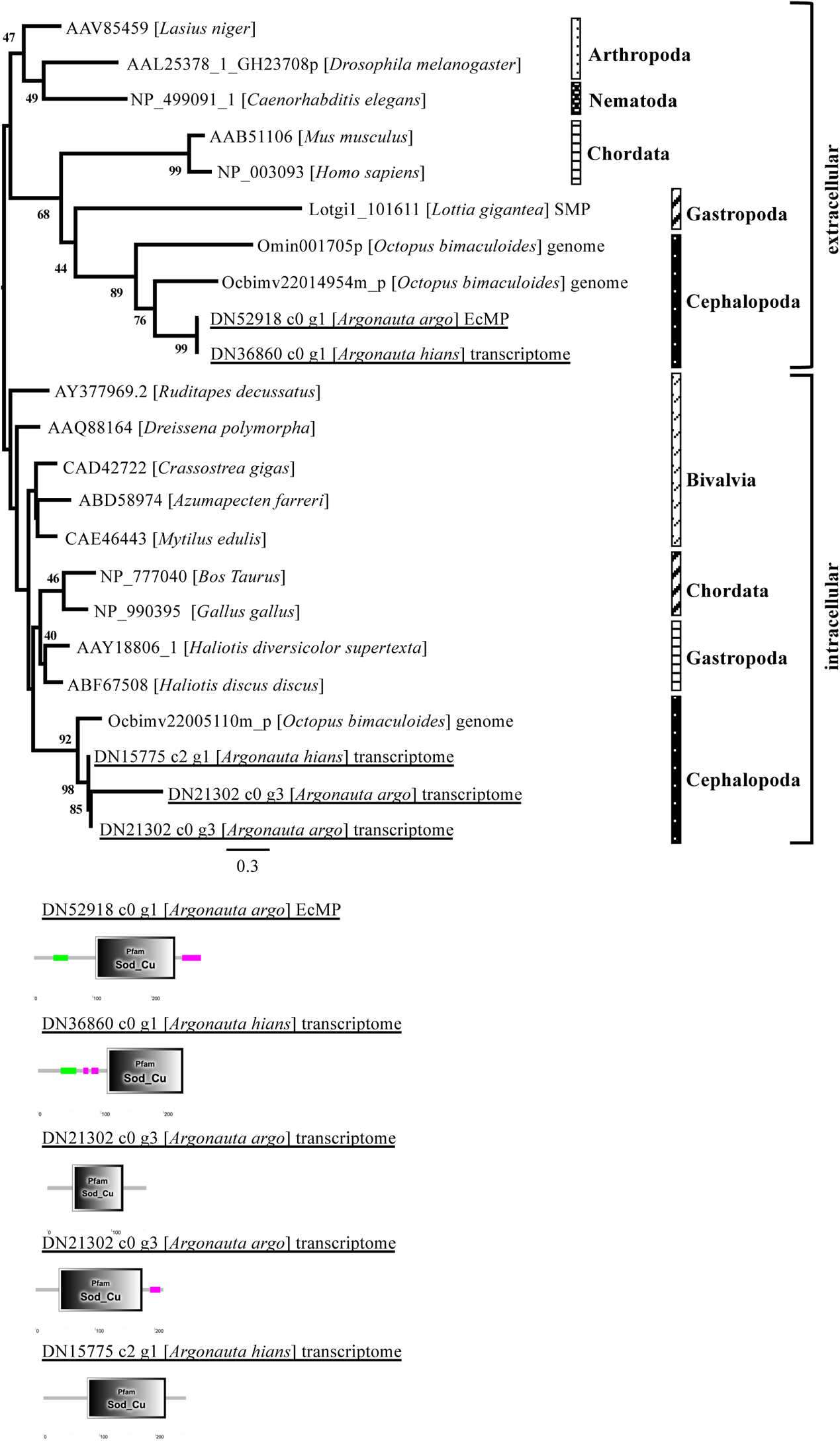
Maximum likelihood phylogenetic trees of selected EcMPs found in the Argonauts. Numbers on the nodes are Bootstrap Support (BS) values. BS lower than 41% are shown as “--”, while 100% support is not written. Representative structures of the proteins of the sequences included in the analyses, shown as SMART protein domains, are shown below the trees. A. Glypicans (Substitution model: LG + Γ + I) B. Thioredoxins (TRX) (Substitution model: LG + Γ + I) C. Pif/Pif-like proteins (Substitution model: WAG + Γ) D. Superoxide Dismutases (SOD) (Substitution model: LG + Γ + I)

Thioredoxin (TRX) is a well-characterized protein (especially in humans), which functions by reducing other proteins through Cysteine thiol-disulfide exchange (Holmgren, 1989). The protein has a single domain, the Thioredoxin (TRX) domain, with the presence of multiple CXXC motifs, which are important for the protein’s function, as the characteristics of the domain. It has been found to be present in all living organisms, and thus, across the whole of Metazoans, including Mollusks. We found TRX in the EcMPs of both species of *Argonauta* targeted in our study. In our phylogenetic tree, the TRX sequences of *Argonauta* formed a monophyletic clade with the sequences of other Cephalopods, including the shell-less ones. The clade was then placed together with the sequences of other Lophothrochozoans (Fig. 5B)

Superoxide Dismutase (SOD) is an antioxidant enzyme, whose function is to facilitate the dismutation of superoxide radicals, catalyzing them to become “ordinary” oxygen molecules. Because of its importance, it is present in all living organisms. SOD is a metalloenzyme, and can be classified into three categories based on its metal cofactors: the Cu-Zn type, the Fe/Mn type, and Ni type. Only the first one is known to be present in eukaryotes, with the Fe/Mn type known to be present only in both prokaryotes and the mitochondria of eukaryotes. Consistent with this classification, we detected the presence of only the Cu-Zn type SOD in both *Argonauta*. Only one sequence of Cu-Zn SOD was found in the EcMP of *Argonauta argo*. Meanwhile, from the transcriptome data of the mantle tissue, three sequences of *A. hians* and one of *A. argo* were detected. In the phylogenetic tree, all of the Metazoan homologs of Cu-Zn SOD were grouped into two monophyletic clades, which is consistent with the classification of Cu-Zn SOD proteins (Ni et al. 2007; Parker et al. 2004): the intracellular type (without signal peptide) and the extracellular type (with signal peptide). Our transcriptome data indicate that the Argonauts secrete both functional proteins in the mantle and the membrane of the first arm pair. Although only one extracellularly secreted protein was detected in the eggshell matrix proteome data of *A. argo*, a homologous sequence was detected in the transcriptome data of the first arm membrane of *A. hians*, and thus, the absence of the extracellular sequence in the EcMPs of *A. hians* were probably artifactual.

Glypican has not been previously identified as a common SMP in Conchifera, thus making our finding interesting. The protein, which is a proteoglycan protein with a single domain, has been implicated in the regulations of various signaling pathways involved in embryonic development, such as Wnt, Hedgehog, and BMP pathways (De Cat et al. 2001; Filmus et al. 2008). The protein has also been reported to be present in the matrix of the chicken eggshell (Mann and Mann 2015). Our phylogenetic analyses showed that the *glypican* gene was probably duplicated in the putative Ur-Conchifera, as attested by the placement of two monophyletic clades of molluscan Glypican sequences together with their Lophotrochozoan homologs (Fig. 5C). The three Glypican sequences found in both *Argonauta* were placed together with their Cephalopod homologs, which then further included within the clade containing Conchiferans, and then Lophothrochozoans (Fig. 5C). It is to be noted, however, that the argonauts’ Glypicans formed a monophyletic clade, and thus probably an autapomorphy of the *Argonauta* clade. This might imply an independent adoption of Glypican in the mineralized eggcase formation of the group.

Tyrosinase, an oxidizing enzyme with a single domain (the Tyrosinase (Tyr) domain), which one of its known functions is to control melanin production in eukaryotes, is also known as a major SMP in Conchiferans. Although the enzyme was not detected in the EcMP proteome of the Argonauts, it was expressed in abundance in the membrane of the first arm and the mantle tissue of both *Argonauta*, as indicated by the transcriptome data. Five were found in *A. argo* and four in *A. hians*. Phylogenetic analysis on a sequence data set containing Tyrosinase homologs in the Cephalopods showed that Tyrosinase sequences were divided into two monophyletic clades, one of which consisted of only the sequences of shell-less octopus, *O. bimaculatus* and the two species of *Argonauta*, one sequence for each species, and the other containing the sequences of all Cephalopods including the *Ilex* squid, the Argonauts, and the shell-less octopuses (Fig. 6)

**Fig. 6.**
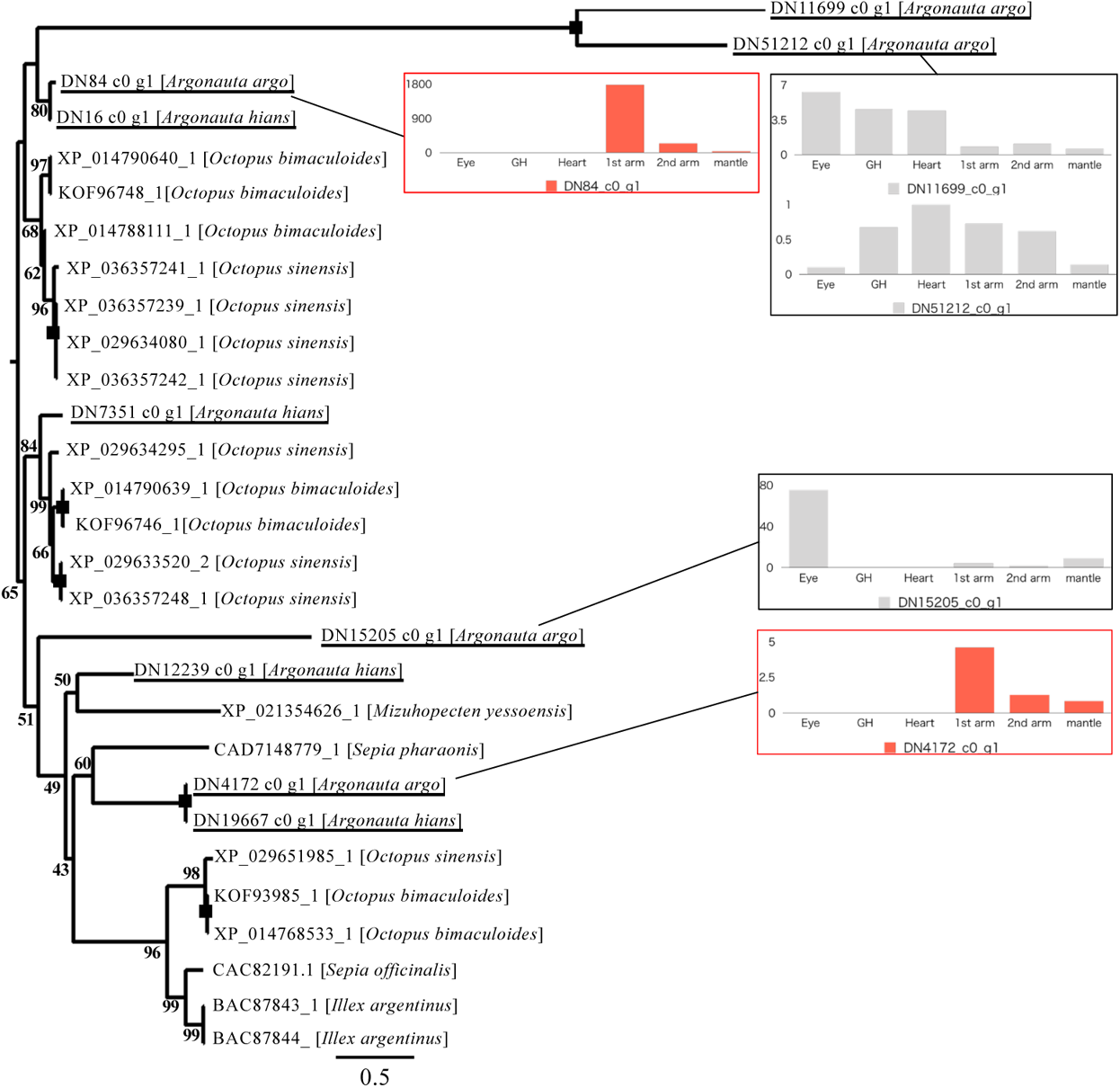
Phylogenetic analysis of Tyrosinase found in the Argonauts. The tree was inferred using Maximum Likelihood inference methods. Numbers on the nodes are Bootstrap Support (BS) values. BS lower than 41% are shown as “--”, while 100% support is not written. Gene expression patterns in *A. argo* are also shown in the figure.

## Discussion

### 1. An Overview of Conchiferan Shell Matrix Proteins

Mineralogy and microstructural studies have shown that the shells are composed mainly of calcium carbonate crystals and consist of several calcified layers (such as prismatic and nacreous) and one organic layer (periostracum). The outer shell of the Conchiferan Mollusks is composed of about 95% calcium carbonate and about 1-5% organic components (Lowenstam and Weiner 1989). Although there are several polymorphs of calcium carbonate crystal (e.g., calcite (trigonal), aragonite (orthorhombic), and vaterite (hexagonal)), the molluscan shells are generally composed of a combination of aragonite and calcite (Spann *et al*. 2010; Frenzel and Harper, 2011). The mineralization process, organization, and maintenance of the molluscan shells are thought to be controlled genetically (Carter 1990; Marin et al. 2012), in a process called “organic matrix-mediated mineralization”, where the formation, growth, and arrangement of biomineralized structures are controlled by biomolecules secreted by the cells, such as proteins, which act as frameworks for the biominerals to precipitate (Lowenstam 1981; Weiner et al. 1983).

A combination of transcriptome and proteome analysis has been used to identify these proteins (e.g., Marie *et al*. 2012; Miyamoto *et al*. 2013; Arivalagan *et al*. 2017; Zhao *et al*. 2018; Setiamarga *et al*. 2020), with some data validated at the genomic level because of the availability of draft genomes (Mann *et al*. 2012; Zhang *et al*. 2012; Takeuchi *et al*. 2016; Kenny *et al*. 2020). Many of these proteins are present in trace amounts inside the shell and are therefore referred to as SMPs. Despite their low abundance in shell matrices, SMPs play an essential role in shell formation and structural maintenance, including calcium carbonate nucleation, crystal growth, and selection of polymorphs of calcium carbonate (Addadi et al. 2006; Marin et al. 2008). Many of these proteins are characterized by composition of both RLCDs and highly conserved enzymes, such as peroxidases, CAs, chitinases, acidic calcium-binding proteins, and protease inhibitors (Marie et al. 2012). Other SMPs include acidic proteins involved in CaCO_3_ crystallization, chitin-binding proteins, and ECM-related proteins (Pif, BMSP, EGF-ZP, SPARC) (Dyachuk, 2018; Feng et al. 2017).

The finding of conserved proteins and/or highly conserved domains across the Conchiferan Mollusks indicates that SMPs have existed as ancestral genes for a long time, but such SMPs have been co-opted independently in different molluscan lineages (Kocot et al. 2016; Setiamarga et al. 2020). For example, independent recruitments of CAs were found in the shells of *C. gigas* and *P. fucata* after their divergence (Zhao et al. 2020). In addition, SMPs classified as RLCDs have evolved rapidly and in parallel (McDougall et al. 2013; Jackson et al. 2010), while Glycine-rich RLCDs, Shematrin, and KRMP gene families have undergone multiple duplications, extensive sequence divergence, and shuffling of various motifs (Yano et al., 2006; McDougall et al. 2013; Kocot et al., 2016). However, the presence of similar SMPs and their domains (although most likely repeatedly recruited and thus not necessarily homologous) and the usage of aragonite and calcite as biominerals have become the trademark of a “true” Conchiferan shell.

### 2. The Argonaut eggcase is not a homolog of the Shell of the Conchiferans

Despite being phylogenetically placed among shell-less Octopodiforms, and thus, a member of the Conchiferans (Hirota et al. 2021), the Argonauts produce a shell-like external structure, which functions as an eggcase, made from calcite (Bøggild 1930; Lowenstam and Weiner 1989), with the outer form resembles the shape of the coiled shells of the Nautiloids and Ammonoids (Scales, 2015; Stevens et al. 2015). This shell-like eggcase is thin and brittle, and unlike the “true” shells of the Conchiferans which were formed by the mantle tissue, the eggcase was reportedly formed by two specialized dorsal arms (Scales, 2015), and thus is thought not to be a homologous structure to the calcified shells of Conchiferans and Cephalopods, but an evolutionary innovation of the genus (Naef, 1923). Moreover, the eggcase is atypical for a Conchiferan shell because it contains chitosan instead of insoluble chitin (Oudot et al. 2020b).

Our study reported here is the first to try to answer the lingering questions of the evolutionary origin of the eggcase, using the multiple omics approach. When summarized, the result is very interesting, because in general, the result is very clear and thus provides a genetic and molecular support to the traditionally accepted notion that the eggcase of the Argonauts is not a homologous structure to the Conchiferan shell. The result is a prime example of the independent adoptions of an array of proteins to produce a converging morphological structure, and thus, although there could be a deep homology in a some of the associated genes, most of the gene repertoires being used to form the mineralized, shell-like eggcase are completely different from those used (and conserved) in the Conchiferan SMPs (Fig. 3 and 4).

### 3. Proteins other than Conchiferan SMPs were Independently Adopted for the Formation of the Argonauts’ Eggcase

#### 3.1. Pif/Pif-like Proteins

The evolution of Pif/Pif-like proteins is complex. Based on their domains, there are at least three groups of Pif/Pif-like proteins: Pif, Blue Mussel Shell Protein (BMSP), and the Laminin G3/Pif-like (Fig. 5A) (Varney *et al*. 2021; This study). Pif was originally found in the nacre of the pearl oyster (Suzuki *et al*. 2009), and *in vitro* functional analysis has shown that it is involved in calcium crystallization (Suzuki *et al*. 2013). In pterid bivalves, Pif mRNA encodes a peptide that is cleaved into two functional proteins, Pif 97 and Pif 80, according to their protein molecular weights (Suzuki *et al*. 2009). Pif 80 is a key acidic protein that regulates the formation of the nacreous layer, while Pif 97 is associated with the polysaccharide matrix of the shell. Domain structures of these proteins also hint at their different roles in biomineralization. For example, Pif 80 binds to pearls and helps them form (Suzuki et al. 2009), while Pif 97 binds to chitin and leads to the crystal growth of calcium carbonate (Suzuki et al. 2013). Proteins with these domain structures are found in a wide range of Mollusks (Suzuki et al. 2013). BMSP, the second homologue of Pif, was detected as SMPs of the blue mussel *Mytilus galloprovincialis* and *L. gigantea*, respectively, although it differs from Pif in domain compositions (Suzuki et al. 2011; Marie et al. 2017). Characterization study by Suzuki et al. (2011) identified that the BMSP is a signal-peptide containing secreted protein, which can bind calcium carbonate crystals of both aragonite and calcite, and thus probably plays an important role in maintaining structure within the shell matrix. The protein also contains multiple VWA domains and one ChtBd, and is present throughout the nacreous layer with dense localisation in the myostracum, and thus suggested its possible role in Conchiferan nacreous layer formation.

We did not detect any of the homologs of Pif and BMSP in the eggcase matrix of the Argonauts. Instead, we found only sequences of the third homolog type of Pif/Pif-like proteins, the Laminin G3/Pif-like, in the EcMP of *Argonauta hians* and in the mantle transcriptome data of *A. argo*. Laminin G3/Pif-like was identified by Marie *et al*. (2017) as one of the two Pif-like isoforms including the following domains; one VWA, three ChtBd, one RLCD or one LamG. Interestingly, the RLCD of the first homolog type of Pif/Pif-like, the Pif proteins, would be identified as a LamG domain in some domain searches, thus suggesting a possible shared origin of the two proteins. Although no extensive functional characterization of the protein has been decisively done, these similarities suggest that Laminin G3/Pif-like could probably be associated with calcium crystallization. Indeed, the protein has also been shown to be associated with biomineralization of shells in the pond snail *Lymnaea stagnalis* (Ishikawa et al. 2020). Since both Pif and Pif-like homologs were found in the chiton, orthologs of both Pif and LamininG3/Pif-like were probably already present in the basal molluscan lineage, suggesting that both homologs were already involved in the formation of a sclerite skeleton (Varney et al. 2021).

Interestingly, no Pif or BMSP could be found in the Cephalopods, and while the homologs of Laminin G3/Pif-like were found throughout the taxon regardless of the presence or absence of a mineralized external shell. In our phylogeny, the Laminin G3/Pif-like proteins of the Argonauts were placed together in a monophyletic clade with other Laminin G3/Pif-like sequences of the nautilus, the octopods, and the bivalves. When considered altogether, we could hypothesize a possible scenario of how the Laminin G3/Pif-like was recruited in the shell formation of the Cephalopods. First, the three types of Pif/Pif-like were probably present (or even used as SMPs) in the putative last common ancestor of the Conchiferans. However, the Cephalopods lost the first two (Pif and BMSP) and opted to use only the third type (Laminin G3/Pif-like). The shell-less octopods still keep the gene Laminin G3/Pif-like for other, presently unknown, reasons. The Argonauts would then re-adopt this gene for the formation of the mineralized eggcase. Thus, this may be an example of an ancient gene that has been independently recruited or lost many times for the formation of the mineralized shells or shell-like mineralized external structures. Meanwhile, the varied copy numbers of Pif/Pif-like proteins in the Conchiferans could probably suggest that independent gene duplications, besides losses, might have occured during the evolution of these proteins.

#### 3.2. Glypican

Glypican, a heparan sulfate proteoglycans implicated in various regulatory processes at the cell surface (Filmus *et al*. 2008), was found in the argonaut EcMPs. We did not find any Glypican in the SMPs of any of the Conchiferans besides Cephalopods surveyed in this study, while its involvement in Conchiferan shell formation had never been reported before. However, interestingly, Glypican is one of the matrix proteins found in the avian eggshells, with one of the isoforms, Glypican-4, found to be shared between eggshell matrices of the turkey and the quail (Mann and Mann, 2015). Together with Osteopontin, eggshell Glypican has a widespread distribution among the vertebrates, and their expressions are induced in eggshell gland epithelia by the mechanical strain exerted upon entry of the egg into the gland (Lavelin *et al*. 2000; Lavelin *et al*. 2002). These inter-phyla similarities strongly suggested that Glypican plays an important role in biomineralization, and thus was independently recruited by the Argonauts for their shell-like biomineralized eggcase formation.

#### 3.3. Absence of Major SMPs in the Cephalopods

Some of the SMPs involved in nacre formation have been identified from comparative studies on several Conchiferan SMPs. Nacrein, a nacre specific carbonic anhydrase that converts carbon dioxide and water into bicarbonate and hydrogen ions, was isolated from *P. fucata* (Miyamoto et al. 1996). The gene for the homologous protein N66 (Kono et al. 2000) was also isolated from *Pinctada maxima*. Mucoperlin was identified from the nacreous layer of *Pinna no*bilis (Marin et al. 2000), and Perlucin and Perlustrin from the nacreous layer of *Haliotis laevigata* (Weiss et al. 2000; Blank et al. 2003). In addition, N16 (Samata et al. 1999) and its homologous gene, Pearlin, were identified from *P. fucata* (Miyashita et al. 2000). Nacrein and N66 are homologous genes in *P. fucata* and *P. maxima*, respectively, and consist of a carbonate dehydrogenase domain and a GN repeat sequence (Miyamoto et al. 1996). A similar homologous gene has also been isolated from *Turbo marmoratus* from a mantle cDNA library (Miyamoto et al. 2003). N16 and Pearlin, isolated from the pearl oyster, belong to the same family, as does N14 from the white-lipped pearl oyster (Kono et al. 2000). Pearlin is a glycoprotein with polysaccharides attached to sulfate groups (Miyashita et al. 2000). There is a report by Samata et al. (1999) on the precipitation of aragonite crystals using N16.

Our studies here suggested that the orthologs of these genes (e.g., Nacrein and Shematrin), which products are crucial for the formation of a mineralized shell with nacreous layers, are not detected in the octopus genome, and also, not in the EcMPs of the Argonauts. Our previous study on the SMPs of the basal Cephalopod *Nautilus pompilius* also did not detect the presence of Nacrein and Shematrin in the shell matrix, although there is a possibility that this could be attributed as an experimental artifact (Setiamarga et al. 2020). When considered altogether, these results could be interpreted as a possibility that the genes were lost in the lineage leading to the Cephalopods, which in the end might lead to their reduced ability to form a mineralized external shell, and then finally culminated in a complete shell loss in the octopods lineage. This could be the reason why the Argonauts invoked a different set of protein repertoires, rather than the traditional SMPs, to form its shell-like eggcase (Fig. 3 and 4). Further studies involving a thorough reanalysis of the nautilus at the genomic level (Zhang et al. 2021) and the Decapodiforms, which still retain some remnants of the shell such as the gladius (e.g., squids) and the cuttlebones (e.g., cuttlefishes) might help to further test this hypothesis decisively.

One of the most abundantly expressed SMPs and widely conserved in the Conchiferans was the protein EGF-ZP, a nacreous layer-specific gene (e.g., Marie et al. 2011b; Marie et al. 2012; Marie et al. 2013; Feng et al. 2017). Generally, this SMP consists of EGF-like domains, which is characterized by six conserved cysteines bound by a characteristic pattern of disulfide bonds and two small beta-sheets, followed by one zona pellucida (ZP) domain. EGF-like domains are present in a variety of proteins associated with diverse biological functions such as cell adhesion, signal transduction, and Ca^2^+ binding. Sequences of the EGF-ZP protein with conservation of each domain (signal peptide, EGF, ZP) were found in Nautilus, suggesting that the genes are probably conserved in the Cephalopods (Setiamarga et al. 2020). Although this protein was not found in the EcMPs of the Argonauts, it was found in the transcriptome of both *Argonauta* species, and in the octopus genomes (e.g., XP_029647132.2 in *O. sinensis*, XP_014769935.1 in *O. bimaculoides*). However, the genes and the protein products were apparently not recruited, and thus not used for the formation of the eggcase. Although further studies are still needed to clarify what would be the effect of losing this gene, and how the expression of the gene is regulated in the formation of the “true” shell of the Conchiferans, it is probably related to the microstructural organizations of the calcium carbonate crystals in the shell matrix. Therefore, the Argonauts probably did not adopt this protein for their eggcase formation, despite its presence in the genome (as attested by its presence in the transcriptome data) because they would need to recruit a complex set of proteins. Because of the absence of EGF-ZP, the eggcase of the Argonauts lacks the nacreous layer, and is composed of only calcite.

However, Oudot et al. (2020a) analyzed the coiled shell of the Spirulid *Spirula spirula*, and found that the species shares only a few similarities with known SMPs of the Mollusks. They also showed the presence of Chitinase, Tyrosinase, and Matrilin, and the absence of RLCD proteins as the characteristics of the Spirulian shell. Meanwhile, the presence of housekeeping genes such as actin and histone, and the absence of EGF-ZP, are apparently shared between *Spirula* and the Argonauts.

### 4. Possible secretory mechanisms of Argonaut EcMPs

It is thought that the shell-like eggcase of the Argonauts is formed by protein secretions from the umbrella membrane of the first arms into the microspace between the umbrella membrane and the eggcase (Scales, 2015). In Mollusks, the space between the mantle and the shell is called the extrapallial space, and it is always filled with extrapallial fluid (Crenshaw 1972). In this study, we also confirmed that the argonaut EcMPs have a secretory signal sequence, including also for Laminin G3/Pif-like and SOD, thus implying that they work in an extracellular secretory manner like the SMPs of the Conchiferans. In addition, the mineral grows through an intracrystalline protein matrix and the membranes between crystal layers in nacre formation (Cartwright and Checa 2007). The nacre assembly shows huge dynamics including the initial chitin crystallization, the subsequent formation of a liquid crystal, its transformation into a protein-coated membrane, and the part played by the extrapallial liquid (Cartwright and Checa 2007). It is conceivable that crystal growth may be very limited in the argonaut eggshells, where chitosan is a major component, consistent with the lack of detection of typical SMPs.

Our multi-omics study reported here without doubt has succeeded in providing a big piece of important information regarding the formation and evolution of the shell-like eggcase of the Argonauts as a convergent morphological structure at the molecular level, which involves independent adoptions of unconventional genes and proteins. Our study also suggests that, in order to understand the eggcase formation of the argonauts, which our analysis indicated are formed by different protein components, it is necessary to assume a quite different mechanism. Therefore, future studies including analyses of extrapallial fluid (e.g., Hattan et al. 2001; Ma et al. 2007), local pH regulation, pigmentation mechanisms, the actual shell formation by using living specimens of the Argonauts that can be reared for short periods of time or individuals that show the recovery process of damaged shells.

## Acknowledgments

Samples of *Argonauta argo* and *A. hians* were obtained as bycatch, presented to us by Minoru Yoshida of Miyuki Suisan, Co., and colleagues in Oki Island. Computations were partially performed on the NIG supercomputer at ROIS National Institute of Genetics (NIG). We are grateful to Kazutoshi Yoshitake and Takeshi Kawashima for their help on how to use the Singularity module on the NIG supercomputer system. We are grateful to the staff researchers of Medical ProteoScope Co., Ltd., for their technical support in proteome analysis. We also thank Kazuyoshi Endo (The University of Tokyo) for his invaluable advice.

This work was supported by the FY2016 Research Grant for Chemistry and Life Sciences from The Asahi Glass Foundation (awarded to DHES) and Grants-in-Aid for Scientific Research (C) No. 18K06363 (awarded to MAY and DHES), and partially supported by the Human Frontier Science Program Grant (RGP0060/2017; awarded to MAY), Nakatsuji Foresight Foundation FY 2018 Research Grants for Basic Sciences (to DHES), Grants-in-Aid for Scientific Research (C) No. 19K12424 (awarded to DHES), and Grants-in-Aid for Exploratory Research No. 19K21646 (awarded to DHES and TS). Some of the transcriptome sequencing was supported by the Platform for Advanced Genome Science (PAGS) Project of Japan (https://www.genome-sci.jp/e). We also thank the faculty of Life and Environmental Science at Shimane University for the financial support in publishing this report.

## Author contributions

MAY and DHES conceived the idea, initiated, and managed the course of the study. HO and NS contributed to the sample collection. MAY and DHES dissected samples, which were then identified and vouchered by TS at The University Museum of The University of Tokyo. HK, RI, MAY, and DHES conducted data analysis. DHES, HK, and MAY conducted molecular works. SI, HY, and YI assisted and advised protein sample preparations conducted by HK and DHES. HK, RI, and MAY conducted bioinformatic analysis. MAY and DHES wrote the first draft of the manuscript, which were then edited further by DHES, HK, TS, and MAY. All authors read, edited, commented, and confirmed the content of the final version of this manuscript.

---

**Supplementary Figure 1. The EcMPs in two Argonauts.**

**Supplementary Figure 2. A comparison among Cephalopoda under search setting 2, shown without the homologs of Zinc-finger proteins.** The result indicates that the number of homologs in the two Argonauts are roughly the same.

