## Supplementary figures and images for "Independent adoptions of a set of proteins found in the matrix of the mineralized shell-like eggcase of Argonaut octopuses"

### Supplementary Figures

#### Supplementary Figure 1

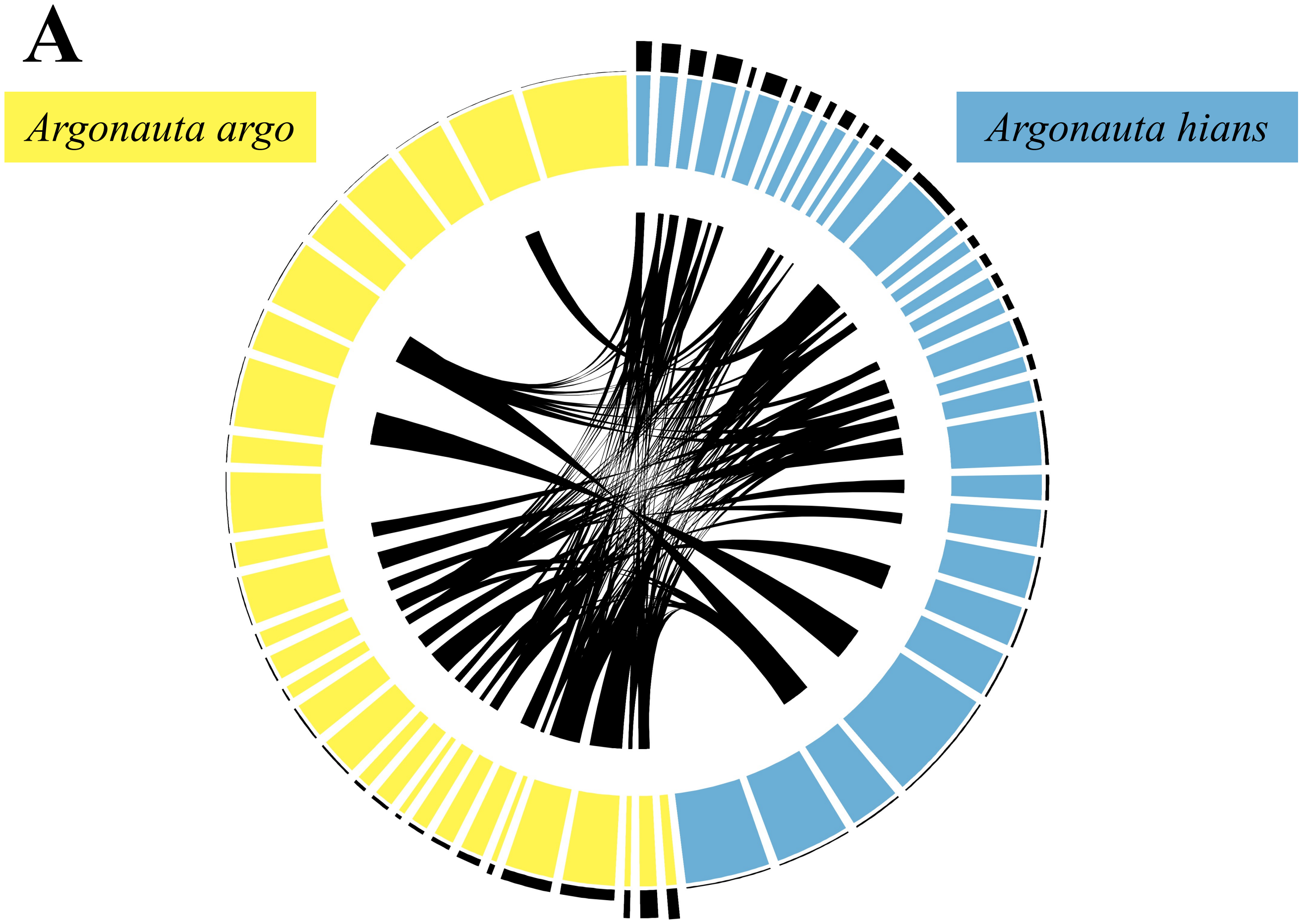


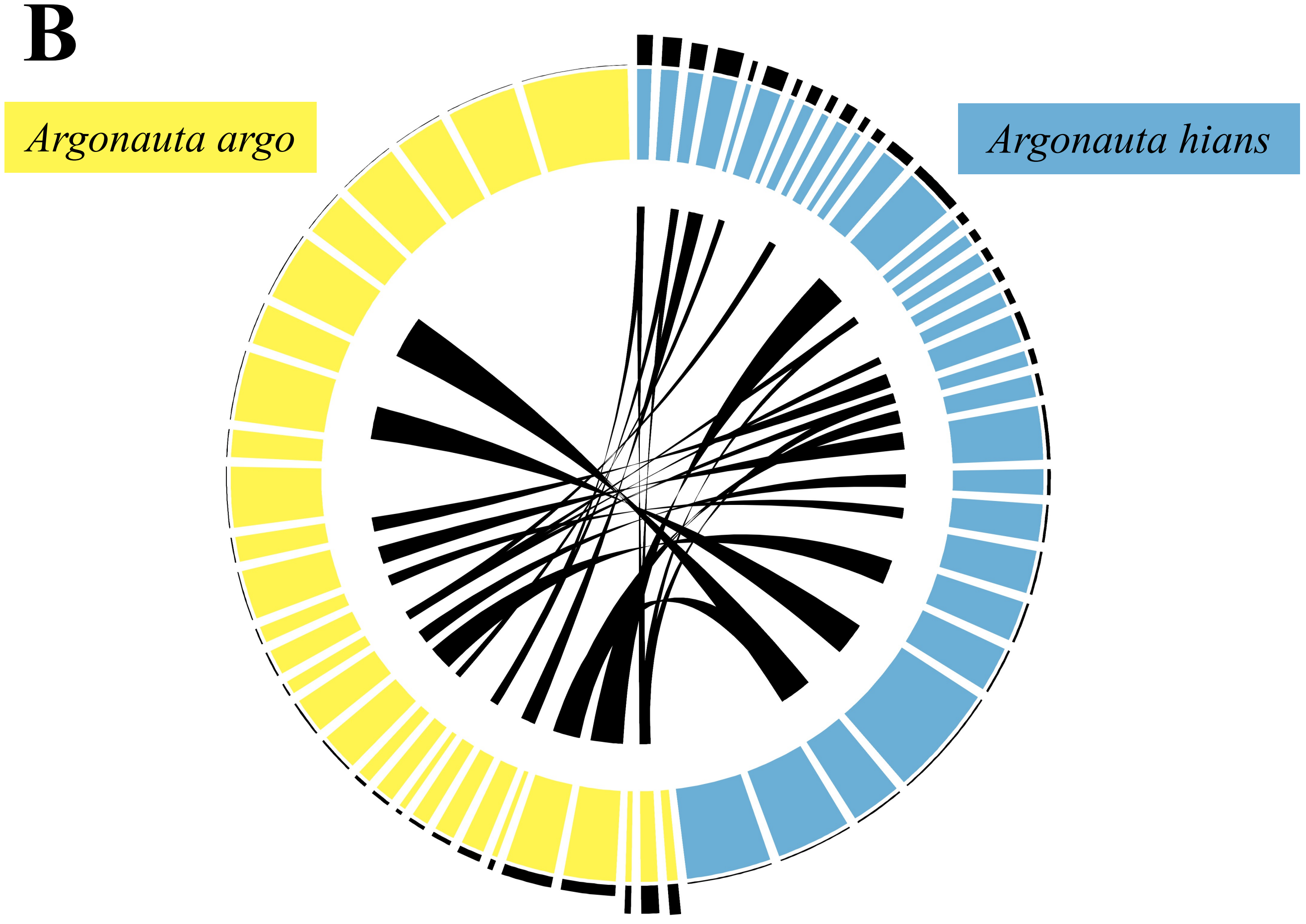

#### Supplementary Figure 2

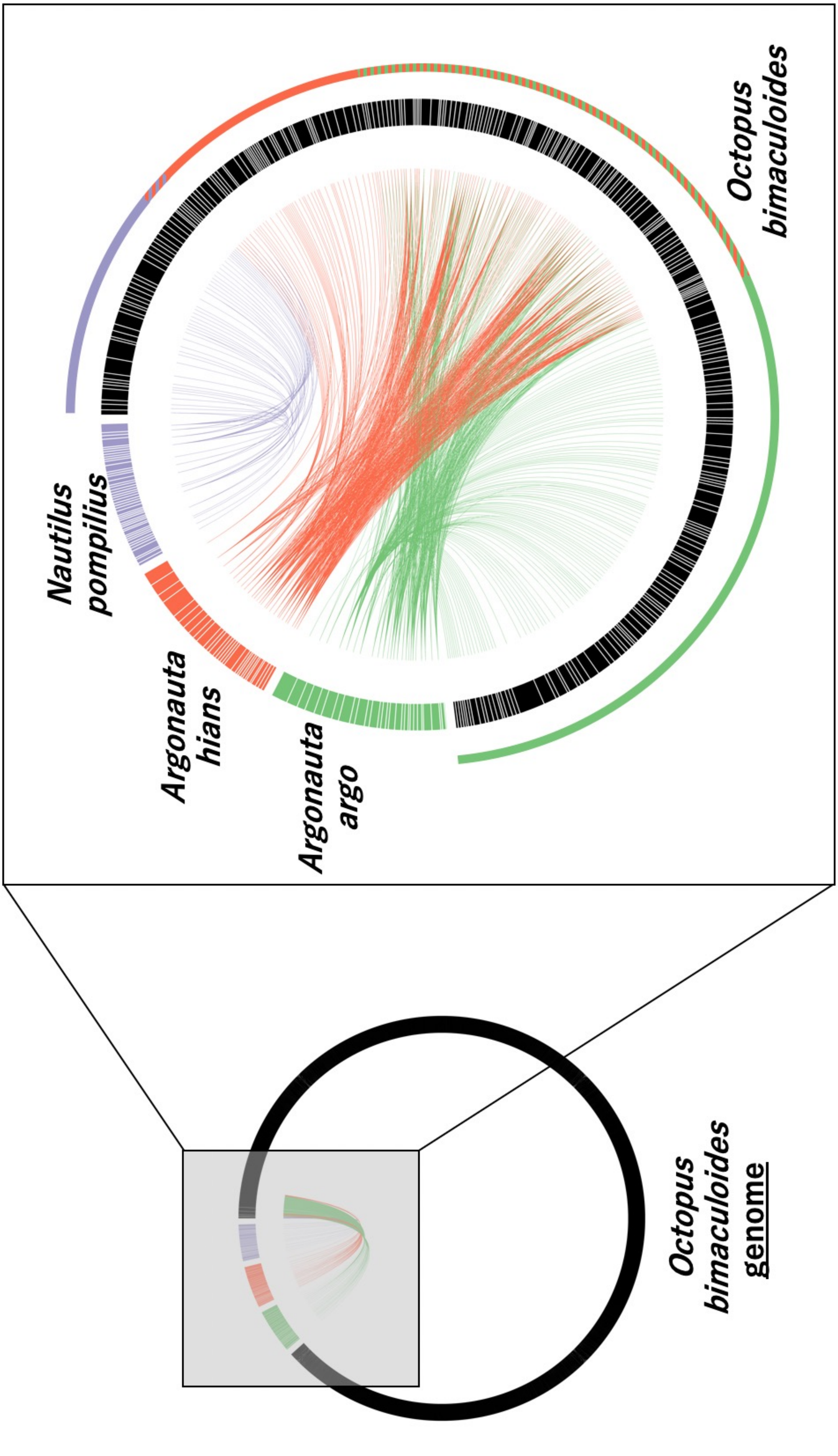
